# Topological damping in an ultrafast giant cell

**DOI:** 10.1101/2021.12.13.472465

**Authors:** Ray Chang, Manu Prakash

**Affiliations:** Department of Bioengineering, Stanford University, Stanford, CA, 94305; Woods Institute for the Environment, Stanford University, Stanford, CA, 94305; Chan Zuckerberg Biohub, San Francisco, CA 94158

**Keywords:** ultrafast biophysics, organelle topology, vertex model

## Abstract

Cellular systems are known to exhibit some of the fastest movements in biology - but little is known as to how single cells can dissipate this energy rapidly and adapt to such large accelerations without disrupting internal architecture. To address this, we investigate *Spirostomum ambiguum* - a giant cell (1-4 mm in length) well-known to exhibit ultrafast contractions (50% of body length) within 5 msec with a peak acceleration of 15_*g*_. Utilizing transmitted electron microscopy (TEM) and confocal imaging, we discover a novel association of rough endoplasmic reticulum (RER) and vacuoles throughout the cell - forming a contiguous fenestrated membrane architecture that topologically entangles these two organelles. A nearly uniform inter-organelle spacing of 60 nm is observed between RER and vacuoles, closely packing the entire cell. Inspired by the entangled organelle structure, we study the mechanical properties of entangled deformable particles using a vertex-based model, with all simulation parameters matching 10 dimensionless numbers to ensure dynamic similarity. We demonstrate how entangled deformable particles respond to external loads by an increased viscosity against squeezing and help preserve spatial relationships. Because this enhanced damping arises from the entanglement of two networks incurring a strain-induced jamming transition at subcritical volume fractions, which is demonstrated through the spatial correlation of velocity direction, we term this phenomenon “topological damping”. Our findings suggest a new mechanical role of RER-vacuolar meshwork as a metamaterial capable of damping an ultra-fast contraction event.

**Significance Statement:** Little is known about how single-cell organisms with extreme motility can decelerate or dissipate energy, as they lack connective tissues. Our study discovered a novel entangled rough endoplasmic reticulum (RER)-vacuolar meshwork architecture in *Spirostomum ambiguum*, an ultrafast giant cell that can contract itself with 15*g* accelerations. We demonstrate through an entangled deformable particle model that the entangled architecture increases the squeeze-flow viscosity of particle systems and helps dampen the motion, a phenomenon we called “topological damping”. For biologists, our study suggests the mechanical role of RER through topological constraints on nearby organelles. For physicists, we point out a new way to create a system with strain-induced jamming. For engineers, we present a novel architecture that can provide braking functions.

---

**H**igh accelerations impart physiological advantages to organisms across all length scales, from speeding cheetahs to the ultra-fast strike of a mantis shrimp(1, 2). In these systems which require repeatable deployment, energy dissipation is critical for the survival of the underlying structures(3). In movement driven by muscles, active lengthening is observed to specifically dissipate energy, and failure to do so can lead to fractures, torn ligaments, or muscle sprains(4). Specialized structures such as foamy mesocarp layer in durian shell (3, 5), and mineralized helicoidal fibrous structures (telson) near the tail of a mantis shrimp (2) are examples where biological structures protect themselves from damages due to impulse. However, how single cells can resist large forces and dissipate energy in short time scales remains largely unknown. Without cell walls or connective tissues, can a single-celled organism evolve strategies to effectively decelerate and dissipate energy as well?

## *Spirostomum ambiguum* as a model system for repeatable ultrafast cellular contraction

*Spirostomum ambiguum* is a giant single-celled organism living in brackish waters, and their extreme contractility is well documented even in early literature(6). (Fig. 1) As a millimeter-scale single-celled ciliate, *Spirostomum* can repetitively contract themselves to less than 50% original body length in 5-10 msec(7) throughout its lifetime. Unlike single-shot ultrafast events such as nematocyst firing (8) which leads to an exploded cell, *Spirostomum ambiguum* demonstrates repeatable contractions via a reversible process of fast contraction and slow relaxation to the base state. The contraction and force generation process is relatively well-known in the literature of *Spirostomum ambiguum*. Briefly, the contraction is caused by the myoneme fibers, an analog to centrin, located in the cortical region. Once you flood the cell with calcium, all the filaments start to contract.(9) In our recent studies, we discovered a novel function associated with ultra-fast contractions in *Spirostomum ambiguum* - to enable long-range hydrodynamic communication(7). The ultra-fast kinematics is necessary to break the Stokesian regime and enable long-distance wave propagation. The kinematics of this process is composed of a rapid acceleration phase immediately followed by rapid deceleration, with peak magnitude of both acceleration and deceleration up to 15*g*(7). Previous studies have also measured the peak contraction force to be about 0.6 *µ*N(10), which is at least 10^4^ times greater than the usual forces experienced at cellular level (roughly 10 pN)(11). A back-of-the-envelope estimate based on the kinematics of the contraction also predicts a strain rate (0.2m/s*/*(1mm*/*2) ∼400sec^*−*1^) that is comparable to the wall shear rate in the large artery of a healthy human(12).

**Fig. 1.**
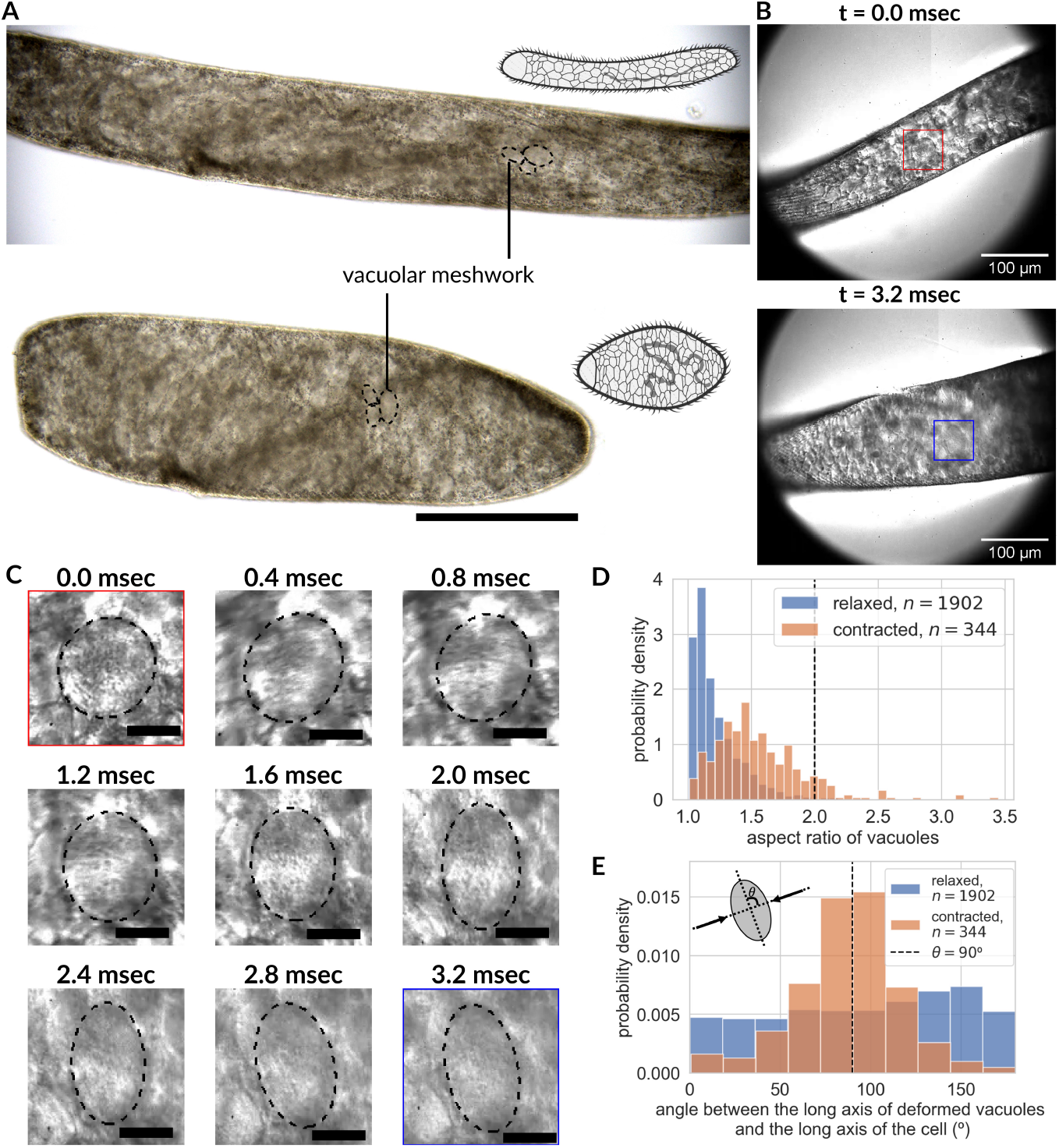
Vacuolar meshwork inside a giant cell *Spirostomum ambiguum* undergoing ultra-fast contractions, leading to organelle deformations. (A) Color differential interference contrast (DIC) images of the same *Spirostomum ambiguum* cell before and after an ultrafast contraction event, depicting ∼50% length shortening in less than 10 msec while maintaining cell volume. The cytoplasm is filled with vacuoles packed at high density. Scale bar 200 *µ*m. (B) Single cell contraction under high-speed camera reveals large-scale deformation (40% deformation from spherical to ellipsoidal) in vacuoles under compressive strain in the first 3.2 milliseconds of the contractile process. The red box marks one vacuole before contraction while the blue box marks the same vacuole after a single contraction event. The image sequence of the detailed shape change under this compressive load for the same vacuole is shown in (C) at a 400-microsecond interval. Also see Movie S1&S2. Scale bar 20 *µ*m. (D) The histogram of the aspect ratio of deforming vacuoles in relaxed and contracted organisms quantitatively establishes that the vacuoles are indeed deformed due to the ultrafast contraction process and relax back to minimum energy spherical shape at a longer time scale. The vertical dashed line indicates 40% deformation. (relaxed: mean = 1.22, std = 0.19, 1902 measurements from 3 organisms; contracted: mean = 1.55, std = 0.35, 344 measurements from 3 organisms.)(E) The histogram of the angle between the long axis of the vacuoles and the long axis of the cell body before and after the contraction depicts the deformation is normal to the compressive stress. Here we define the direction of the long axis of the cell body as the *x*-direction, and the direction of the short axis as the *y*-direction. (relaxed: mean = 96.1^*°*^), std = 50.8^*°*^, 1902 measurements from 3 organisms; contracted: mean = 87.6^*°*^, std = 28.9^*°*^, 344 measurements from 3 organisms.

## Existing mechanisms fail to explain the energy dissipation in *Spirostomum*

Despite this high strain rate, we can quickly do a back-of-the-envelope calculation to demonstrate that fluid shear alone is still not enough to dissipate the energy injected into the system. Assuming that the input power from cortical myoneme is balanced by fluid shear alone, we can write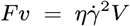, where *F, v, η*, 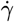 and *V* are boundary force, velocity, cytoplasm viscosity, shear rate, and cytoplasm volume, respectively. Because of the slender cylindrical shape of the cell, we can estimate its volume as *LD*^2^, where *L* and *D* are the length and diameter of the cells. Plugging in the relevant numbers (*F* ∼ 10^*−*6^N, *L* ∼ 10^*−*3^m, *D* ∼ 10^*−*4^m, *v* ∼ 0.2 m/s, 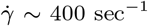 sec^*−*1^), it shows that the viscosity of the cytoplasm has to be as large as 0.12 Pa-sec to dissipate the input energy, which is 120 times larger than the viscosity of the water. This number is also in direct contradiction to the previously measured cytoplasm viscosity in other ciliates or protists, which is about 0.005-0.05 Pa-sec(13, 14). These contradictions suggest that it is not possible to dissipate the energy just by the fluid shear in the cytoplasm if the cytoplasm were completely filled with liquid without structures. Also, the myoneme itself also cannot provide enough brake for itself. In the past, Ishida has performed ghosting experiments (9, 15), in which the cortical myonemes are preserved but the internal structures are disrupted by detergents. If you increase calcium in those ghost cells, the entire cell will collapse and crumble. This indicates that there is not enough intrinsic brake in the myoneme itself. At the same time, we would expect such a high strain rate in the cytoplasm can cause massive rearrangement of organelles, which can potentially break the inter-organelle contact sites which are critical for the physiology. How can then *Spirostomum ambiguum* evolve a mechanism to resist such a high force and decelerate effectively while maintaining the internal architecture?

One unique feature that has been noticed in *Spirostomum ambiguum* for years is their highly vacuolated cytoplasm (also known as vacuolar meshwork, with the diameter of individual vacuole ranging from 5 *µ*m to 30 *µ*m) (Fig. 1A & SI Appendix, Fig. S9)(16), which is very uncommon except for some other giant single-celled ciliates such as *Bursaria truncatella, Homalozoon vermiculare*, and *Loxodes magnus*(17). In giant phytoplanktons, vacuolation is believed to increase the surface area:cytoplasm ratio and maximize resource acquisition(18). In mammalian cells and even in protists, cytoplasmic vacuolization is usually associated with various pathological conditions and is often accompanied by cell death(19, 20). However, this is clearly not the case for *Spirostomum ambiguum*, and the biomolecular function of these vacuoles remains poorly understood. By imaging live cells undergoing contraction at high frame rates, we discovered that individual vacuoles deform up to 40% (see Method for estimation of strain) within the cytoplasm akin to a deforming shock-absorbing foam (Fig. 1B-C, Movie S1-S2). Is it possible that the highly vacuolated cytoplasm actually plays a critical role in decelerating the contraction process and participates in energy dissipation?

## Discovery of entangled architecture between RER and vacuoles

In order to investigate energy dissipation in this unique cell and identify any damage that might be caused to specific organelles - we map key organelles inside a cell utilizing high resolution imaging such as confocal microscopy and transmission electron microscopy (TEM). Several organelles like the rough endoplasmic reticulum (RER) and mitochondria are distributed across a cell and are good candidates to map. RER particularly, not only forms contiguous networks throughout the cells(21), but also makes membrane contact sites with other organelles such as mitochondria, Golgi apparatus, lysosomes, peroxisomes, endosomes, and plasma membranes(22). Maintaining the contiguous geometry and membrane contact sites is crucial to their functions in calcium homeostasis, lipid synthesis, lipid exchange, and protein secretion.(22)

To understand how ultra-fast cells can protect the organelle architecture and inter-organelle contact sites while undergoing massive accelerations and deceleration - we decided to map the RER topology with respect to vacuole present in *Spirostomum ambiguum*. To our surprise, we discovered a fenestrated-cubic membrane-like topology of RER, which is closely wrapping around the vacuolar meshwork. (Fig. 2A) Under TEM, the distribution of vacuoles is consistent with our observation in light microscope and early histology studies, with a thin layer of periplasm (2-3 *µ*m) near the cortex enclosing a highly vacuolated endoplasm which spans through the entire organism(16, 23, 24) (Fig. 2B & SI Appendix, Fig. S1). The intimate wrapping relationship we describe here between RER and vacuoles is preserved in both relaxed and contracted organisms (Fig. 2A). This unique topology (wrapping) is present throughout the entire cell (Fig. 2B). When quantified across the cell - the distance between RER and associated vacuole also has a tight distribution between 30-70 nm in relaxed organisms, suggesting stereotypical membrane contact sites (MCS) between these two organelles. In contracted organisms, the distance between the RER and its nearby vacuoles has a slightly wider distribution, but it still falls below 150 nm (Fig. 2B inset).

**Fig. 2.**
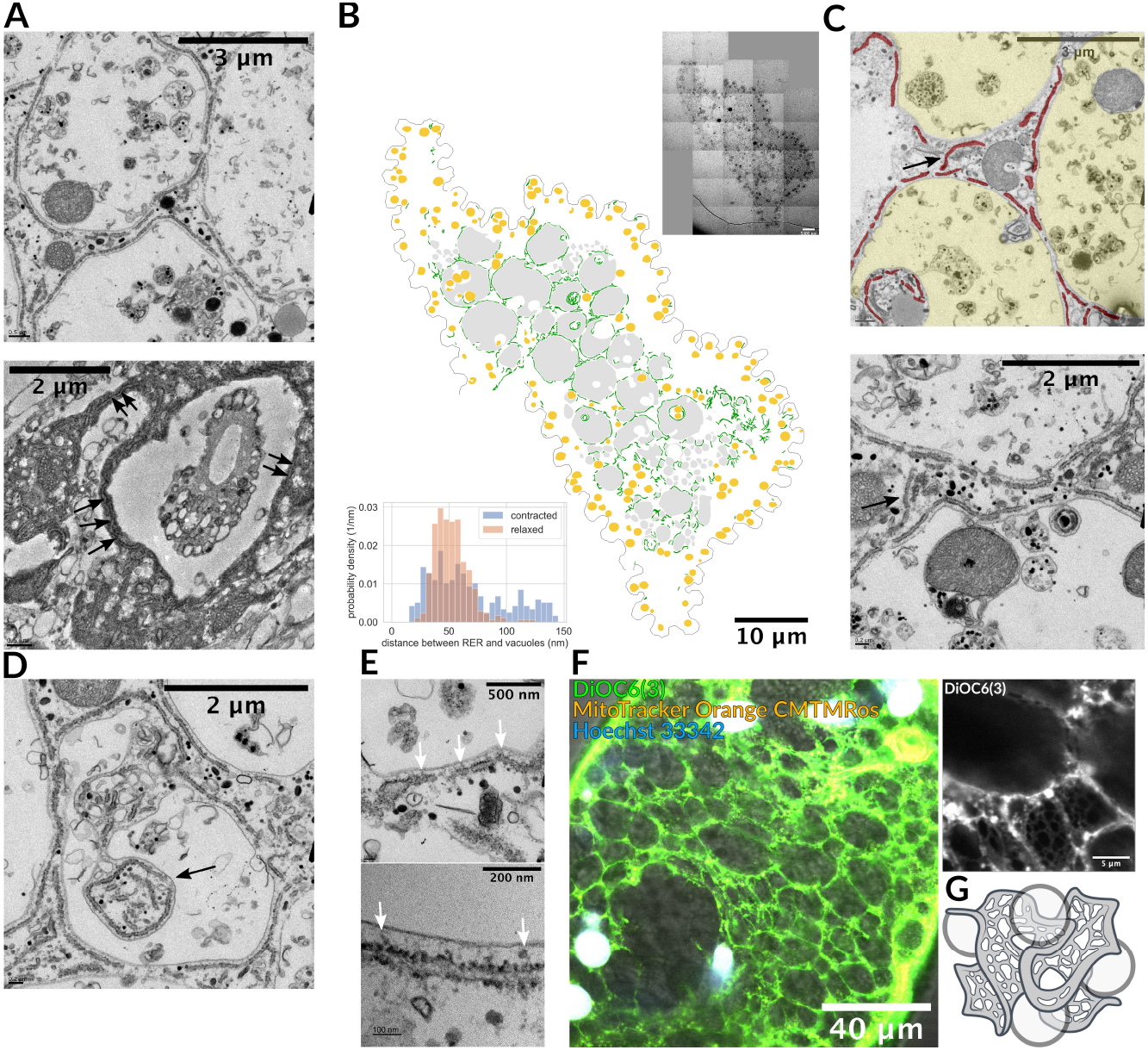
Intimate spatial relationship between rough endoplasmic reticulum (RER) and vacuolar meshwork. (A) TEM imaging of *Spirostomum ambiguum* before (upper) and after (lower) ultrafast contraction shows the vacuolated cytoplasm, and the RER is intimately wrapping around the vacuoles in both cases. Refer to SI Appendix, Figure S1 for low magnification images. (B) Cross-section montage of a relaxed organism and its segmentation of RER (green), vacuoles (gray) and mitochondria (yellow). The entangled topology between RER and vacuoles spans the entire cross-section, not limited to a specific region. The inset on the upper right corner shows the original image. Width of original image: 65.287*µ*m. The histogram on the lower left corner shows the distance between RER and nearby vacuoles in contracted and relaxed organisms under TEM. In the relaxed organism, the distance ranges from 30 to 70 nm (343 measurements, mean±std: 51.9±14.6nm), while in the contracted organism, this distance becomes more variable but remains below 150nm (269 measurements, mean±std: 82.1±52.9nm). (C) TEM occasionally reveals the “bridging” RER (arrows) that connects the RER of two nearby vacuoles. This is expected as RER is a continuous lumen. RER false colored in red and vacuoles in yellow on the top figure. (D) Extreme tram-tracking of RER along the contour of vacuoles was observed, even when cytoplasm invaginated inward to the vacuoles (arrows). (E) At higher magnification, we can see some materials (white arrows) that might be the protein connections between RER and vacuoles. (F) Confocal imaging of endoplasmic reticulum (ER) and vacuolar meshwork of a contracted organism (left) and a cropped magnified region (upper right). Note that the image is a longitudinal section. The image shows a fenestrated web-like structure of ER wrapping around vacuolar meshwork throughout the entire organism. Refer to Movie S3-S5 for complete z-stack video. (G) 3D schematic drawing illustrating the entangled topology between fenestrated ER and vacuoles.

Since RER network is contiguous throughout the cell - we also directly observe bridging RER that travels from one vacuole to another nearby vacuole, which is expected for forming a continuous lumen in the cell (Fig. 2C). We also find that the RER is fenestrated, with holes puncturing the membrane into many segments. Even when the cytoplasm is invaginated into the vacuoles, the RER still closely follows the outline of the vacuoles (Fig. 2D). All the above evidence suggests that there is a linker between RER and vacuoles in *Spirostomum ambiguum*, either through individual proteins or physical forces like Casimir effect (see discussion for more detail). Although we cannot clearly identify the mechanism of this linker as yet, we did observe some structures under high magnification which might be associated with proteins between RER and vacuoles (Fig. 2E). A more careful structural study is required to identify their identity.

To observe the 3D architecture of the RER-vacuole interaction, we image fixed organisms under confocal microscopy, with the staining of DiOC6(3), an ER marker(25) (Fig. 2F, SI Appendix Fig. S2, Movie S3-S5). The finding in confocal microscopy confirms our observation under TEM. We clearly see a fenestrated sheet wrapping around the vacuoles throughout the entire cytoplasm, and the highly deformed vacuoles (as the organisms fixed for confocal microscopy were all contracted) seem caged within the similarly deformed RER. All these observations confirm the presence of a unique metamaterial composed of RER topologically intertwined with the vacuolar meshwork - throughout the cytoplasm (Fig. 2G). The extremely narrow gap between RER and vacuoles we observed under TEM, the fact that RER closely follows the contour of the vacuole even when the cytoplasm is invaginated, and the similar deformation between RER and vacuoles under confocal microscopy all indicate that RER and vacuolar meshwork are mechanically coupled with each other.

## Role of entanglement in bulk material properties

Our data illustrate a unique geometry of entangled RER and vacuolar meshwork inside a contracting cell. Next, we explore the physiological consequences of this geometry for this ultra-fast contracting cell. Previous studies have shown that highly entangled systems can impart unique mechanical properties, both in living and nonliving matter. Some examples of single-component entangled objects include fire ants - which are known to interlock themselves via limbs and mandibles, and thus form waterproof rafts and towers that can be crucial for their survival after floods(26). In simulations of tissues, inclusions can cause marked stiffening of the extracellular matrix, which might explain the compression stiffening of various biological tissues(27). Even simple geometries like entangled U-shaped particles like staples can stabilize the structure against vibration through inhibition of particle rotation and translation(28). In polymer rheology, topological characteristics of entanglement, such as linking number and writhe, have been shown to be related to their mechanical properties like storage modulus and loss modulus(29). Entangled polymer rings, compared to their linear counterparts, also have less shear thinning behavior(30). In two-phase entangled materials - engineered wire-reinforced granular columns have been shown to build freestanding stable structures with high strength under vertical load.(31) Recently, interpenetrating lattices have shown enhanced damping properties in 3D printed materials.(32)

## Entangled soft particle vertex model

Inspired by the geometrical arrangement of RER and vacuolar meshwork, we seek to find if such a system can act as a damper and control the kinematics of contraction at sub-cellular length scales. In the study of granular material, 2D models have been shown to be very informative to explore the fundamental mechanisms and are used extensively when new constraints such as particle deformability or surface frictions are introduced(33, 34). The current model captures the dynamics of entangled RER and deformable vacuoles in a contracting, area-preserving geometry. As we are trying to isolate the effect of entangled topology, we currently focus our efforts on this coupling between these two organelles instead of the detailed cellular hydraulics(35).

We first build a 2D entangled soft particle model representation of this unique metamaterial to study its mechanical properties when the system experiences extreme shape changes (Fig. 3; SI Appendix, Fig. S3), using a vertex-based model with smooth surface method(33). In our model, the cell is simplified into a two-dimensional rectangle, the vacuoles in the cytoplasm are simplified into deformable particles, and the RER is simplified into flexible strings connecting the particles. The topological links introduced by the tethered strings mimic the membrane bridges we found under the TEM, which allows us to explore the role of entanglement in the mechanical properties of this unique architecture(36). (See SI Appendix, Section A.7.1.)

**Fig. 3.**
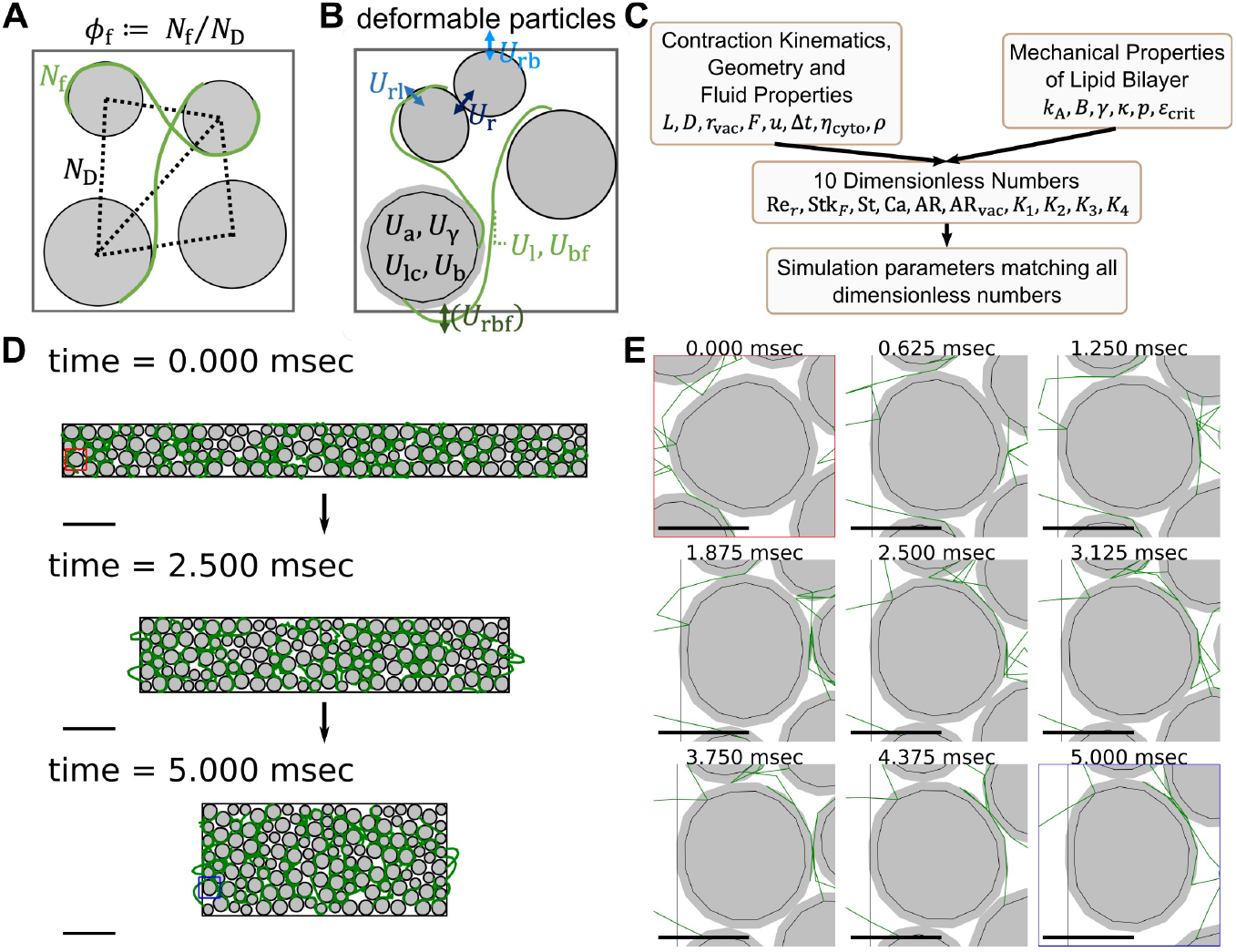
A 2D vertex-based entangled soft particle model allows us to study the effect of topological constraints on the mechanical properties of packed vacuoles in response to large shape changes. (A) We define filament fraction (*ϕ*_f_) as the ratio between the number of strings (*N*_f_) and the number of edges in the Delaunay diagram (*N*_D_), which we use to describe the degree of topological constraints in the system. The volume fraction (*ϕ*) is defined as the ratio between the total area of particles and the area enclosed by the boundary. See SI Appendix, Section A.7.2. (B) We consider 10 energy terms in the model following the recommendation from Boromand et al.(33) with appropriate extensions to include the entangled strings. The system evolves according to an over-damped molecular dynamic scheme. See SI Appendix, Fig. S3. (C) We curated 10 dimensionless numbers from the known contraction kinematics, geometry, fluid properties, and mechanical properties of lipid bilayer membranes, and all simulation parameters are determined to match all 10 dimensionless numbers. The process ensures dynamic similarity and allows us to convert simulation results back to actual physical units and compare them with experiments. (D) Three snapshots of a simulated contraction process. The system displayed here has *ϕ* = 0.79 and *ϕ*_f_ = 0.268. We report this particular simulation as it is the one which matches the experimental observation the best (see Fig. 4). A constant boundary force is applied to deform the system during the contraction phase, with *ϕ* kept constant. The red box marks one particle before contraction while the blue box marks the same particle after contraction. The image sequence of the shape changes under this load for the sample particle is shown in (E) at a 625-microsecond time interval. Scale bar = 100 *µ*m in (D), while scale bar = 20 *µ*m in (E). Also see Movie S6-7.

We model vacuoles as a 50:50 mixture of bidisperse deformable particles with a 1:1.4 ratio and the RER as flexible Hookean strings initialized to be on the opposite side of two neighboring particles to ensure entanglement. Based on the number of particles (*N*_c_) and the number of strings connecting the particles (*N*_f_), we also defined filament fraction (*ϕ*_f_) to describe the degree of connectivity or topological constraint in the system, which is defined as the ratio between the number of strings and the number of edges in the Delaunay diagram of the center of the particles (Fig. 3A). Volume fraction (*ϕ*) is defined as the ratio between the total area of particles and the area enclosed by the boundary. The strings have zero thickness, which ensures that any observable changes come from a pure topological effect. (See SI Appendix, Section A.7.1-A.7.2.)

We consider 10 energy terms following the recommendation from Boromand et al.(33), with proper extension to include entangled strings: 1) particle-particle repulsive energy (*U*_r_), representing the repulsion between vacuoles; 2) particle-string repulsive energy (*U*_rl_), representing the repulsion between RER and vacuoles; 3) particle-boundary repulsive energy (*U*_rb_), representing the repulsion between vacuoles and cell membrane; 4) stretching energy of strings (*U*_l_), representing the energy when RER is stretched; 5) bending energy of strings (*U*_bf_), representing the energy when RER is bent; 6) particle surface contractility (*U*_lc_), representing the energy when vacuole membrane is stretched; 7) particle compressibility (*U*_a_), representing the energy when the vacuole area is compressed; 8) particle interfacial tension (*U*_*γ*_), representing the interfacial energy between the vacuolar membrane and cytoplasm; 9) particle surface bending (*U*_b_), represent the energy when vacuolar membranes experience bending. An additional 10th term, string-boundary repulsive energy (*U*_rbf_), is included only during the initialization (see SI Appendix, Section A.7.6). We obtain forces on particles by taking gradients with respect to the coordinates of each vertex, and the system evolves according to an overdamped molecular dynamics scheme. (See SI Appendix, Section A.7.3.)

The computation of the energy terms requires many parameters (e.g., the bending coefficient 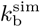, area compressibility coefficient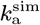, etc. See SI Appendix, Fig. S3 for all parameters). To ensure our parameter choice accurately captures the physics, we next perform dimensionless matching to ensure dynamic similarity. We define 10 relevant dimensionless numbers (Fig. 3B; SI Appendix, Section A.7.4, A.7.11, and Fig. S3) based on known contraction kinematics and known mechanical properties of biological components like lipid bilayer membranes and vacuoles. These include 1) Reynolds number, 2) Stokes number, 3) Strouhal number, 4) capillary number, 5-6) two aspect ratios, and 7-10) four relative ratios among the five mechanical properties of membranes and vacuoles (area expansion modulus, water bulk modulus, interfacial tension, bending modulus, hydration repulsion pressure) (Fig. 3C; SI Appendix, Section A.7.4, A.7.11, and Fig. S3). This dimensionless matching process allows us to convert all output from simulations back to actual physical units and compare them directly to experimental observation.

We contract the system with conserved area for 5 msec (the duration is dimensionally matched to the correct kinematics of cellular contraction) in the long axis (defined as the *x*-axis), imposing fixed boundary forces (magnitude dimensionally matched to the experimentally observed force magnitude(10)) and observe the changes in kinematics (Fig. 3D). The boundary force treatment we used here is a reasonable approximation, as it is well-described in past literature that the organism contracts when calcium binds to cortical myoneme(37), limiting the force to the cortical region. Also, even though we only explicitly set the boundary forces in the *x*-direction, because of the area conservation constraints in the 2D model, there will always be associated movement on the boundary in the *y*-direction, so the effective consequence is a squeeze strain and not compressive strain. Our system allows us to follow the deformation of each particle just like in the experiments (SI Appendix, Fig. S4), and we show the time series of a specific particle in Figure 3E. In the model proposed, the reactive forces created by the particle system counteract the boundary forces, which directly reduce the amount of net energy input (see SI Appendix, Section A.7.5 for its calculation). The net energy input can be (1) buffered by the various potential energies in the system, (2) viscous dissipation by the particle motion within the fluid, or (3) dissipated by the sacrificial bond when the string ruptures. Note that because of the dimensionless matching process, the system has a huge timescale separation (see SI Appendix, Section A.7.5). This requires a small time step even with implicit methods to ensure accuracy. Depending on *ϕ*_f_, it takes 30-60 days to finish one simulation when using 6 CPUs with 4GB memory per CPU. The relaxation phase is not simulated as the deceleration happens during the contraction phase (see Section “Limitations of the model”).

## Damping effect from topological constraints in entangled soft-particle systems

Here we explore how topological constraints impact the kinematics, boundary forces, squeeze flow viscosity, and energetics of the contraction process (see Movie S6-S7 for the simulation videos). In order to understand the role of entanglement, we ran 16 simulations with the same *ϕ* of 0.67 and 0.79 (both chosen to be sub-critical volume fractions for 2D circular disks, below 0.83) but varying *ϕ*_f_. We monitor the time series changes of different observables (Fig. 4) including normalized body length, normalized contraction speed, normalized reactive force, squeeze flow viscosity, and change in the energy budget. In the main text, we only report the results of *ϕ* = 0.79 as the results are similar. See SI Appendix, Figure S5 for the results of *ϕ* = 0.67, and Figure S11 for results with parameters variation.

**Fig. 4.**
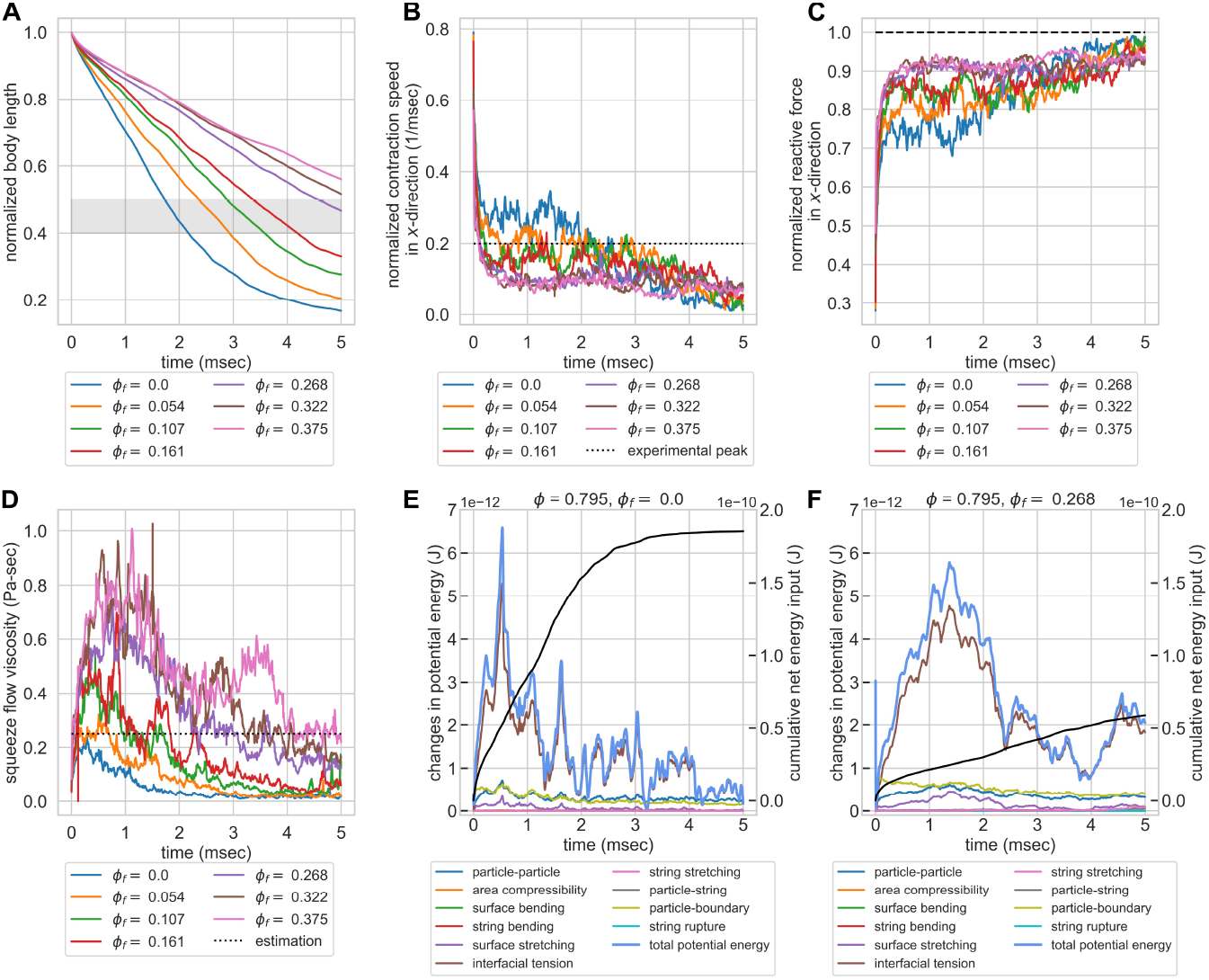
Geometric constraints turn entangled particle systems into topological dampers in constant force simulation. Systems with the same *ϕ* of 0.79 but varying *ϕ*_f_ were compressed by a constant boundary force. (A) Normalized body length over time. The gray zone indicates the experimentally observed final body length after contraction (40%-50%)(7). The system with *ϕ*_f_ = 0.268 falls within the range. (B) Normalized contraction rate in *x*-direction over time. The dashed line indicates the experimentally observed peak velocity (0.2 m/sec) (7). Systems with *ϕ*_f_ ≥ 0.268 can rapidly dampen the velocity, while systems with lower *ϕ*_f_ have their contraction velocity at peak level for roughly 3 msec. (C) Normalized reactive forces in *x*-direction over time. The dashed line indicates the magnitude of external load, normalized to 1. Increasing topological constraints increases the ability of the system to counteract external forces, which explains the rapid slowdown in contraction kinematics. (D) Squeeze flow viscosity of the system as a function of time. The dotted line is the estimation of the actual organism based on the peak strain rate and normal stress in *x*-direction (0.25 Pa-sec, see SI Appendix, Section A.7.8). Systems with *ϕ*_f_ of 0.268 and 0.322 maintain comparable squeeze flow viscosities to experimental estimations, while systems with *ϕ*_f_ *<* 0.268 fail to maintain their viscosities despite the initial peak. (E)&(F) Energy budget of 2 systems with the same *ϕ* of 0.79 but *ϕ*_f_ of 0.0 and 0.268. The cumulative net energy input is plotted in the black line, while other colored lines indicate different energy terms. The system with no entanglement has a larger cumulative net energy input. It rapidly responds to the energy input but is not able to store the energy for greater than 2 msec. Systems with entanglement reduce the cumulative net energy input to one-third and also have a better ability to buffer the energy input. Note that the majority of the energy contribution comes from the particle interfacial tension term, and the contributions from strings or string rupture are minimal.

Since the model parameters are chosen to ensure dynamic similarity, we can compare our outputs to experiments. We first analyze the cell length changes during the contraction process. In Figure 4A, all systems contract monotonically as expected, and systems with greater entanglement finished up at a larger normalized body length after 5 msec. The system with *ϕ*_f_ = 0 ends up at a normalized body length of less than 20%, which is far below the experimentally observed final length (marked by the gray zone in the figure), indicating that the vacuoles alone at a subcritical volume fraction are not sufficient to dampen the kinetics. For our simulation setting, the system with *ϕ*_f_ = 0.268 has a final body length matching the experimental observations.

We also compare the contraction speed in simulation to the experimentally observed contraction speed. In Figure 4B, we can see that systems with *ϕ*_f_ ≥ 0.268 rapidly dampen the contraction speed to be below the experimentally observed peak velocity, while systems with *ϕ*_f_ *<* 0.268 fail to dampen the velocity and maintain their contraction speed at peak level for roughly 2-3 msec, which is unrealistic. This indicates that the entangled topology can act as a damper and slow down the contraction kinematics. Note that the initial peak is much greater than the experimentally observed peak velocity, likely because we did not consider the initial time required to build up cytoplasmic calcium concentration and build up the force to the peak.

As for the reactive force on the boundaries, Figure 4C shows that systems with *ϕ*_f_ *<* 0.268 have a lower force magnitude in general, and the time series of reactive forces show more fluctuation. On the other hand, systems with *ϕ*_f_ ≥ 0.268 have persistently larger force magnitude, up to 90% of the external load, and the time series show less fluctuation. These differences in force magnitude explain the observed differences in contraction speed across different systems and the rapid slowdown in contraction kinematics for systems with a larger degree of entanglement. The high degree of fluctuation in reactive forces for systems with lower *ϕ*_f_ probably comes from the frequent rearrangement of particles and relaxation of forces, while the entangled topology in systems with high *ϕ*_f_ restricts the rearrangement and prevents force relaxation.

Informed by these changes in kinematics and boundary forces, we seek to quantify the changes in mechanical properties of the system and compare that to estimation from experiments. Here we define squeeze flow viscosity (*η*_*s*_) as the ratio between the normal stress in *x*-direction (*σ*_*xx*_) and the contracting strain rate in *x*-direction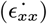, or

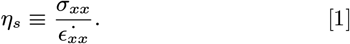

The dotted line is the estimation of the actual organism based on the peak strain rate and normal stress in *x*-direction (0.25 Pa-sec, see SI Appendix, Section A.7.8). Systems with *ϕ*_f_ ≥ 0.268 maintain their squeeze flow viscosity at a level comparable to estimation from experiments, while systems with *ϕ*_f_ *<* 0.268 fail to maintain their squeeze flow viscosity despite the initial peak. This result provides a solution to the energy dissipation conundrum we mentioned previously. Although the cytoplasm viscosity we set for the simulation is only 0.005 Pa-sec (see SI Appendix, Section A.7.4), the entangled architecture increases the squeeze flow viscosity, an effective viscosity of the entire structure, to a value matching our back-of-the-envelope estimation. This increased squeeze flow viscosity in entangled deformable particles supports that the entangled cytoplasmic structure can act as a topological damper for the contraction process.

Our dimensionless-matched model also allows us to dissect the energy budget from different energy terms during the contraction process. In Figure 4E&F, we compare the energy budget of 2 systems with the same *ϕ* of 0.79 but *ϕ*_f_ of 0.0 and 0.268. The system with no entanglement has a larger cumulative net energy input, as the system is less capable of counteracting the forces and there is more displacement. The non-entangled system can rapidly respond to the energy input. However, the system is not able to store the energy as it rapidly relaxes through particle rearrangement. Systems with entanglement can reduce the cumulative net energy input and also has a better ability to buffer the energy input. Note that the majority of the energy contribution comes from the interfacial tension term of the particles, indicating that the majority of potential energy in the system comes from the deformation of the vacuoles. The contribution from string stretching, string bending, and the sacrificial bond from string rupture are all minimal, indicating that the effect of increased squeeze flow viscosity and better ability to hold energy is purely a topological effect (rather than from the mechanical strength of the strings). In SI Appendix, Section A.7.11, we also discuss how variations in model parameters can change the dominant energy terms in the systems. However, even considering the variations in parameters, the energy contributions from string stretching, string bending, and string rupture remain minimal. This suggests that in the real organism, RER does not have to be unrealistically stiff to play a mechanical role, and it can potentially thicken the cytoplasm by its entangled topology with vacuoles alone.

Aside from deceleration, preservation of neighboring relationships is also physiologically relevant, as previous studies have demonstrated the importance of organelle connectomes in other eukaryotic organisms(38). In our simulation, by tracing the neighboring relationships among particles before and after contraction, we can see that the system with no topological constraints shows a massive rearrangement of neighboring particles, which can potentially break the inter-organelle connections if happen in real organisms. On the contrary, the systems with high topological constraints can better preserve the spatial relationship among particles (Fig. 5A&B), suggesting a better capability for entangled organelle architecture to preserve the organelle spatial relationship. This is also consistent with our analysis of the energy budget. As systems with entanglement have less cumulative net energy input and more energy buffered by the potential energy, less amount of energy is needed to be viscously dissipated at each time, which reduces the fluid shear, prevents rearrangement, and preserves neighboring relationships.

**Fig. 5.**
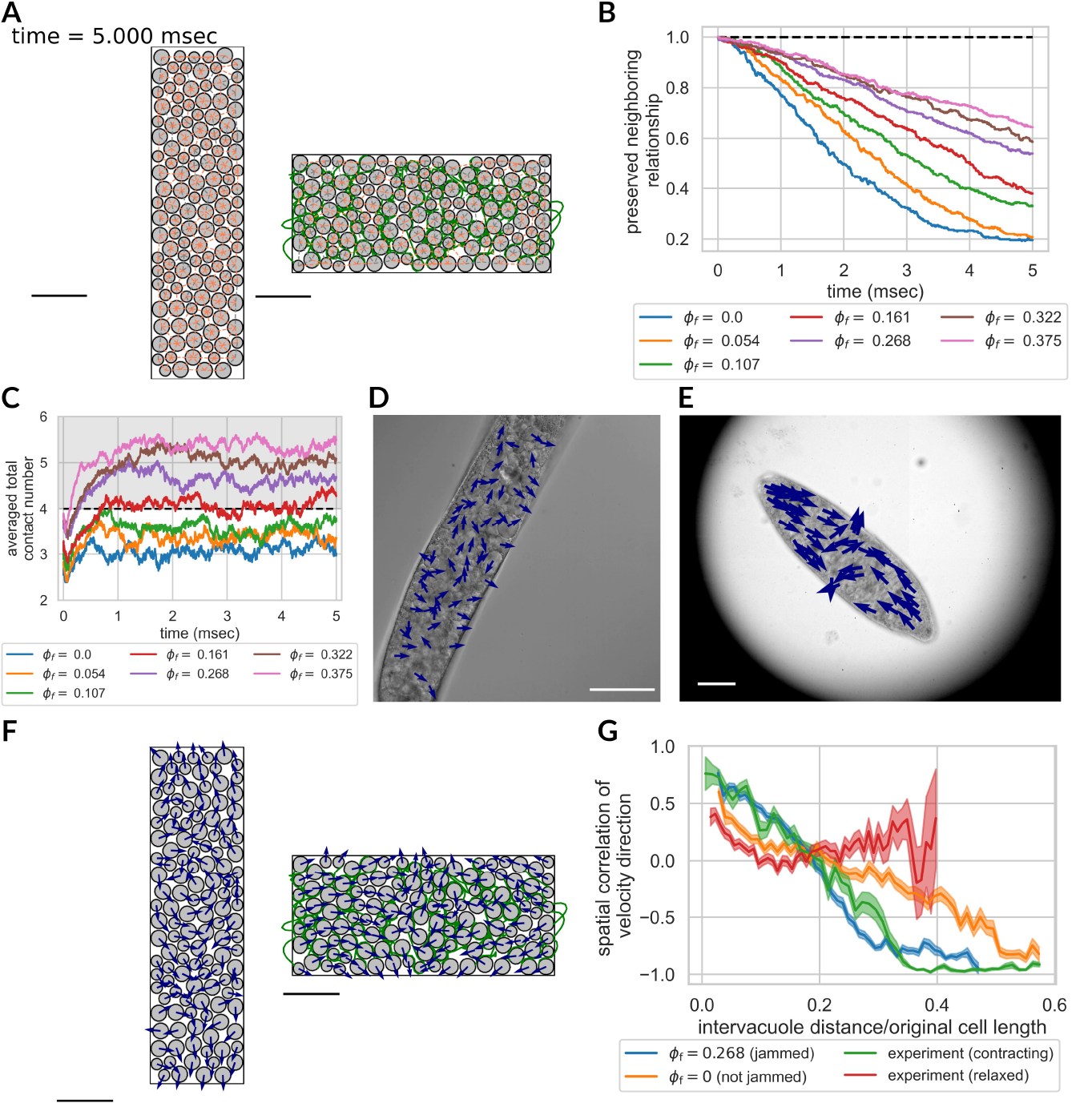
The phenomenon of topologically-assisted strain-induced jamming. The entangled strings create more contact with neighboring particles, making the particles overly-constrained and jammed more easily. (A) Preserved neighboring relationship after contraction for two systems. The neighbors are identified as the edges on the Delaunay diagram. Newly formed neighbors are pink while preserved neighbors are gray. The left figure (*ϕ*_f_ = 0) preserves 19.6% of neighbors, while the right figure (*ϕ*_f_ = 0.268) preserves 54.0%. *ϕ* = 0.79 for both systems. Scale bar = 100 *µ*m. (B) The preserved neighboring relationship of the system over time. Increasing topological constraints helps preserve the neighboring relationship after contraction. (C) Averaged total contact number of the particles over time for systems with different *ϕ*_f_. Systems with *ϕ*_f_ ≥ 0.161 undergo strain-induced jamming transition as the averaged total contact number is above 4(39). (D)&(E) Velocity unit vectors of vacuoles in relaxed (D) and contracting (E) organisms. In relaxed organisms, vacuole movements are disorganized, indicating poor spatial correlation. In the contracting organisms, the vacuoles on both ends are directed toward the center, and the vacuoles near the middle are directed away from the center in perpendicular direction, showing a strong spatial correlation. Scale bar = 100 *µ*m for both. (F) Velocity unit vectors of particles for the same system as in (A) at the end of contraction. The velocity unit vectors for the system with no entanglement (left) show less spatial correlation, while the ones for the system with entanglement (right) show more spatial correlation. Scale bar = 100 *µ*m. (G) Spatial correlation of velocity direction as a function of normalized intervacuole distance (mean ± standard error). Jammed cases in simulations and contracting organisms (3 organisms, total 118 vacuoles) exhibit slower decay in spatial correlation, with negative correlation when normalized distance exceeds 0.3 due to confinement. The unjammed case shows a faster decay in spatial correlation and a negative correlation at larger distances. Relaxed organisms (3 organisms, total 182 vacuoles) show fast decay in spatial correlation at short distances and nearly-zero correlation as the normalized distance goes beyond 0.1. These indicate that the vacuolar meshwork is not jammed in relaxed organisms but jammed in contracting organisms, constituting a strain-induced jamming transition. (Normalized distance where spatial correlation = 0.25 (mean±se): *ϕ*_f_ = 0.268: 0.148±0.006; *ϕ*_f_ = 0: 0.071±0.007; experiment(contracting): 0.147±0.013; experiment(relaxed): 0.040±0.007.) (See SI Appendix, Section A.9, Section C, and Figure S12.)

## Strings thicken the deformable particle systems through topologically-assisted strain-induced jamming transition

From a mechanistic point of view, the phenomenon of topological damping can be explained via strain-induced jamming transition below the classical critical volume fraction threshold due to entanglement. It is well-established that, for 2D hard friction-less spherical particles, when the averaged contact number (⟨*z*⟩) of the particles exceeds 4, the system is jammed because the total number of constraints (*N*_c_ ⟨*z*⟩ */*2) exceeds the total degree of freedom (2*N*_c_) of the system (condition of isostaticity) (39). In our current model for topologically induced damping, we explore how increasing the number of interconnecting strings in the system creates additional contacts between strings and particles under contraction, which also impose additional constraints on the particles and make it easier to be in a jammed state.

Figure 5C shows the averaged total contact number of particles (including both contact from other particles or strings) with respect to time for the same systems in Figure 4. None of the systems plotted started from a jammed state even including the effect of strings. Yet systems with *ϕ*_f_ ≥ 0.268 rapidly enter a contracting strain-induced jamming when the systems start contracting. The system with *ϕ*_f_ = 0.161 is right on the boundary of the jamming transition, while systems with *ϕ*_f_ *<* 0.161 do not jam. Note that as the shape deformation in our simulation is area-preserving, we did not change *ϕ* of the system. The systems have strain-induced jamming at a sub-critical volume fraction due to topological constraints, which prevent the particles from rearrangement and fully utilizing the volume available in the system. These differences in jamming also explain the qualitative differences in behaviors shown in Figure 4, as systems with *ϕ*_f_ ≥ 0.268 show a very different pattern in reactive forces and squeeze flow viscosity.

To further quantitatively validate this effect, we provide a prediction on the mean contact number contributed from strings using probability theory. For points distributed according to a Poisson process in the plane with constant intensity, the average neighbor of each vertex on the Delaunay diagram is six(40). *ϕ*_f_ of the system can thus be estimated as *ϕ*_f_ ∼ *N*_f_*/*(6*N*_c_*/*2). The total perimeter of the particles in the system is 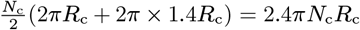, where *R*_c_ is the baseline radius (see SI Appendix, Section A.7.2). As defined in SI Appendix, Section A.7.2, the natural length of each string is 1.4× mLf × *R*_c_. The total length of strings in the system can then be estimated as (1.4mLf*R*_c_)(3*N*_c_*ϕ*_f_). As each string is capable of touching 2 particles on each side, the predicted additional contact from strings (⟨*z*_f_⟩ _predicted_) can be predicted as

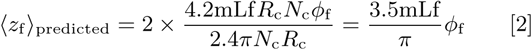

In SI Appendix, Figure S7 we compare the ratio between the observed mean contact number from strings in simulations (⟨*z*_f_⟩) to our prediction, and we see that the time trajectory of the ratio ⟨*z*_f_⟩*/* ⟨*z*_f_⟩ _predicted_ for systems with different *ϕ*_f_ collapse to the same curve and reach a plateau of 1.2. This provides quantitative guidance for how we can produce strain-induced jamming systems with topological constraints.

## The vacuolar meshwork of *Spirostomum ambiguum* undergoes strain-induced jamming transition

Experimentally, however, it is very hard to quantify the contact number of vacuoles from microscopy images even if we have the highest resolution in 3D as RER and cytoplasm occupy the space between neighboring vacuoles. We thus use the spatial correlation of velocity direction as a surrogate(41) to demonstrate the strain-induced jamming transition in *Spirostomum*. Figure 5D and E show the velocity unit vectors of vacuoles in relaxed and contracting organisms, respectively.

The vacuole movements in relaxed organisms come from the cytoplasmic streaming. The velocity direction spatial correlation in contracting organisms is much stronger than in relaxed organisms, showing a clear pattern that vacuoles on both ends are directed toward the center, and vacuoles near the middle are directed away from the center. As we compare these observations with the known jammed and unjammed cases in simulations (Fig. 5F), we see the velocity directions of vacuoles in unjammed case (*ϕ*_f_ = 0, left) is also less directed, with vacuoles seemingly moving in swirls. As for the known jammed case (*ϕ*_f_ = 0.268, right), the vacuoles on both ends are directed toward the center, and the vacuoles near the middle are directed away from the center in a perpendicular direction, similar to the observation in contracting organisms in Figure 5E. Quantitative measurements of velocity direction spatial correlation confirm these observations (Fig. 5G). The spatial correlation of velocity direction in contracting organisms shows a similar decaying pattern as the simulation with *ϕ*_f_ = 0.268 at *t* = 5 msec (entangled and jammed). On the contrary, the spatial correlation of velocity direction in relaxed organisms shows poor correlation even for short intervacuole distances, similar to the pattern from the simulation with *ϕ*_f_ = 0 (not entangled and not jammed). The negative correlations at large intervacuole distances are expected for the two simulations and contracting organisms, as the systems are confined and have boundary movements. The velocity direction spatial correlation patterns indicate the vacuolar meshwork is not jammed in relaxed organisms but jammed in contracting organisms, constituting a strain-induced jamming transition. (See SI Appendix, Section A.9 and Section C.)

## Table-top experiments demonstrate the concept of topological damping

To conceptually demonstrate the idea of topological damping and its potential application and build intuition for this phenomenon, we also created a table-top 3D model of simplified metamaterial capturing the key aspects of this unique ER-vacuole system with entangled table tennis balls and inextensible fabric (Fig. 6, see SI Appendix, section A.8). (Note that here we use hard particles as a demo.) The two systems that we tested have the same number of table tennis balls and the same amount of fabrics, but one includes fabric cut into stripes and entangled with the table tennis ball, while the fabric in the other system simply encloses the table tennis balls in a sealed bag. We can see that the system with entangled topology has a better ability to resist external load (see Movie S8-S9), and the differences can also be seen more quantitatively using compression testing (Fig. 6C, Movie S10-S11).

**Fig. 6.**
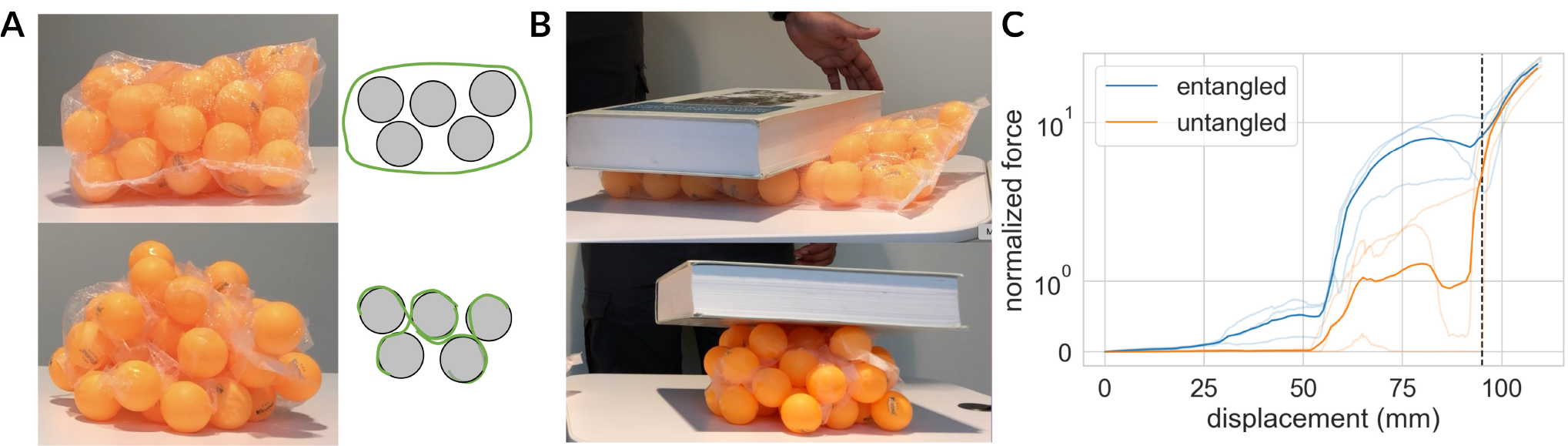
We used 45 table tennis balls and fabrics to demonstrate the concept of topologically-assisted strain-induced jamming. Two systems were compared, with an equal number of table tennis balls and fabrics. One system was arranged in an entangled topology (lower in (A)), while the other was not (upper in (A)) (see Method for details). (B) After applying a 2450 g external load, the entangled system showed superior resistance to the load. Refer to Movie S8 & S9. (C) Quantitative differences between the entangled and untangled systems are demonstrated through compression testing. Mean curves of three samples in each group are shown as solid lines, while individual experimental data are transparently displayed. The system’s reactive forces are normalized by the reactive forces of an individual table tennis ball when compressed by 5 mm (mean ± standard error = 152.1 ± 0.7 N). At a compression of 95 mm (dashed vertical line), corresponding to the onset of unavoidable table tennis ball deformation, a rapid increase in reactive forces was observed in all traces. Refer to Movie S10 & S11.

## Discussion

### Discovery of the novel RER-vacuolar meshwork entangled architecture

In this study, we demonstrate that there is a clear topological association between RER and vacuoles in *Spirostomum ambiguum*. This is the first time that this special topology between RER and vacuoles has ever been observed, even in *Spirostomum ambiguum*. Past TEM data on *Spirostomum ambiguum* either focus on the cortical region(9, 37, 42–49) or focus on other organelles(50–54). Some of the papers mentioned ER but did not correlate the structure with the vacuolar meshwork, most likely due to poor preservation of the vacuolar meshwork(48, 51, 55). For the first time, by performing high resolution TEM and confocal microscopy, we provided extensive evidence of this new topology of the cell, which is preserved in both contracted and relaxed states. Organelles have a very specific spatial relationship with each other, and that is crucial for the fundamental function of the cells. Without the shock absorber, incredible damage can be done to the cell because the inter-organelle connectivity and proximity will be broken. As a single *Spirostomum* would go through at least 300-400 contractions in their life cycle (1 contraction event every 15 minutes without any perturbation, and its generation time is roughly 4 days)(56, 57), the importance of the topological damper on preserving cellular architecture is obvious.

### Mechanical role of RER

Our study also shows that RER can have a mechanical role inside a cell by providing topological constraints on the vacuolar meshwork and inducing jamming transition. In fact, there are some other examples showing the mechanical roles of ER inside the cell. In plant cells, the ER-plasma membrane contact sites have been shown to provide mechanical stability to the plant cells (58). In the Drosophila embryo, intact ER networks are critical to maintaining the pulling forces of the mitotic spindle(59). ER has also been shown to impact the viscosity of cytoplasm, and failure to reorganize ER during mitosis can cause chromosome mis-segregation(60). Our study, however, is distinct from these examples in that the mechanical role of ER network is established in the context of entanglement with other organelles, and the ultrafast nature of the contraction process in *Spirostomum*. Our physically matched model suggests that RER does not have to be unrealistically stiff to thicken the cytoplasm, as in our simulations, the strings bear minimal forces compared to the interfacial tension force of the particles, storing roughly 4% of the potential energy only (for the system with *ϕ* = 0.79 and *ϕ*_f_ = 0.268). In our simulation, we allow the strings to rupture. However, the sacrificial bond of string rupture also accounts for minimal energy compared to particle deformation. The implication that RER can potentially transform the mechanical properties of cytoplasm, not by its own mechanical strength, but instead by creating topological constraints on nearby organelles, warrants us to label this unique entangled architecture as a metamaterial.

### The nature of RER-vacuole contact sites

Although the exact nature of the contact sites between RER and vacuoles is unknown and beyond the scope of this paper, some preliminary results using BLAST (basic local alignment search tool) on the recent transcriptomics data of *Spirostomum ambiguum*(61) showed that there is an expression of proteins analogous to MOSPD2 in *Spirostomum*, a protein that connects RER and lipid droplet(62). (It should be noted though, that the vacuoles are not lipid droplets. See SI Appendix, Section D.) The close and uniform distance between RER and vacuoles also suggests that physical forces such as Casimir effect can be at play. In the past, Bradonjic has discussed how Casimir effect can be the driving force of rouleaux formation in red blood cells, and we can estimate the stable distance between two membranes by differentiating the potential with respect to separation distance(63). Assuming a membrane dielectric constant of 6 (64), and a surface charge density of ER to be 60 mC/m^2^(65), we can estimate the separation distance between the two to be roughly 30 nm, which matches the observed distance in orders of magnitude. As the surface charge density of a lipid bilayer membrane can vary greatly based on the surface protein and lipid composition(65), it is thus possible to hold ER and vacuoles together simply by physical forces. More detailed imaging with antibody staining and measurements is needed to further delineate the nature of this new contact site.

### Evaluations of alternative possibilities

In this study, we focus on the role of organelle architecture in dampening the extreme motility of *Spirostomum ambiguum*. Another obvious candidate for this role is the cytoskeletal network. For example, in the hypodermal syncytia of *C. elegans*, the ER network is anchored to the cytoskeletal network (microtubules and actins) by ANC-1, an ortholog of Nesprin-1/2, and helps maintain the proper positioning of other organelles when the worm is crawling.(66) It is interesting to see that ER again is used for intracellular structural support, though through its anchoring on the cytoskeletal network. In *Spirostomum*, the cytoskeletal network is mostly localized in the cortex. An actin-like protein is found only localized around the macronucleus and along the membranelles, not throughout the cytoplasm(67). Microtubules are only localized in the longitudinal grooves of the cortex(9, 68). Other major structural proteins, such as centrins, spasmins, GSBP1 and GSBP2, are the major players in calcium-triggered ultrafast contraction in *Spirostomum*, but are all localized in the cortex as well(68). None of these are entangled with the ER or vacuolar meshwork in *Spirostomum*. We acknowledge that we did not successfully perform perturbation experiments to demonstrate that RER-vacuolar meshwork is the only dominant factor in maintaining the architecture (see SI Appendix, Section B for our attempts to use ER drugs to perturb ER network), and it is still possible that the ER-vacuolar meshwork is anchored to the cortical cytoskeleton. However, we use our physically-matched model to demonstrate that the entangled architecture is mechanically capable of dampening the motion. Given the spatial separation between the cortical cytoskeleton and other organelles in *Spirostomum*, it is unlikely for just periphery anchoring to cause the jamming transition in the entire cytoplasm without the entangled architecture, even if the anchoring between ER and cytoskeleton exists.

### Topologically-assisted strain-induced jamming

From a physicist perspective, our study also points to a new way to induce jamming(69). In the traditional phase diagram, a granular system can become jammed through cooling, increasing volume fraction, or decreasing external load. In our athermal topologically entangled system, however, the system becomes jammed by increasing external load without changes in volume fraction. For the first time in our study, we described the physics of strain-induced jamming of topologically connected particles. Through velocity direction spatial correlation, we also demonstrated strain-induced jamming transition in an intracellular context. With a physically matched model, we further showed the potential biological role of intracellular strain-induced jamming transition in an ultrafast single-cell organism. A similar phenomenon of strain-induced jamming at a sub-critical volume fraction has been observed in granular particles with frictions(70).

However, the topological constraints we added in the paper are fundamentally different from friction. Firstly, frictions are always limited to nearest-neighbor interaction, while the topological constraints we have can connect and restrain the movement of particles that are beyond the nearest neighbor. Secondly, frictions can only affect two particles at a time, while the topological constraints created by one string can affect more than two particles. Another example that is similar to our study is the jammed wire-reinforced granular column.(31) However, our study is also different from those systems, as the purpose of flexible strings in our model is not to confine free-standing piles into a rigid structure, but instead to convert confined granular materials into an effective damper with other biologically relevant functions.

### Comparisons with other damping materials

From an engineering perspective, it is very interesting to compare the geometry of our system with previously identified structures that are effective in providing braking, shock absorption, or mechanical support. Many types of geometries and architectures have been identified as effective energy absorbers and have wide applications in aerospace, military, and civil engineering(3). These include layered cylindrical structure(71), multi-cell tubes(72), honeycombs(73), foams(74), interdigitated structures(75), auxetic materials(76), and interpenetrating lattices(32). Our table-top experiment demonstrates the potential of our topologically entangled granular system to be a new class of shock-absorbing materials, which still preserve flexibility and permit deformation but can be readily jammed. This provides mechanical strength when experiencing external load and still remains flexible.

### Limitations of the model

To the best of our knowledge, our model is the first model that ever simulates a dense array of deformable biological particles while trying to match all physically relevant dimensionless numbers. Our rigorous simulation approach from first principles allows our model to give explicit predictions on the system with no tuning knobs. Several limitations remain for the model we used despite our greatest effort to make it physically accurate. First, the length of the strings is set to be uniform, while in reality, we would expect the length of bridging ER should have some distribution, even though we do not think this will change the main conclusions of the model. Second, several key features of the contraction process of *Spirostomum* are not accounted for. These include the asymmetry of the contraction process(77) and the torsional component of the contraction process(15). While we expect the asymmetry should do little with respect to the braking of cytoplasm, it is possible for the torsional component to bring the system under topological confinement faster than pure contraction. However, as it is not possible to introduce torsion in a 2D simulation, we will leave that as future work. Third, our model also did not account for the time required to develop boundary forces. It is unlikely for the organism to reach maximum force at time 0 as it takes time to build up the calcium concentration within the cytoplasm. Compared to the actual time series of contraction speed in *Spirostomum*(7), the peak velocity did not occur at time 0, and this discrepancy likely comes from this difference. However, we believe the general conclusion of our paper should still hold, as that simply changes the amount of energy input in the system and does not impact the energy dissipation mechanism inside the system. Also since the detailed time development of the boundary force in *Spirostomum* is not known, our model presents the most simple case with no additional assumptions. Fourth, our model did not account for all possible interactions between RER and vacuoles and only focused on the topological effects. From our observations in Figure 2 and our previous discussion, it is very likely that the interactions between RER and vacuoles include attractive forces, in addition to the effect from entangled topology. This discrepancy might explain why the observed particle deformation is not as large as the deformation in actual vacuoles. Fifth, in our model, we did not simulate the relaxation process. As our paper focuses more on energy dissipation and damping of contraction kinematics, it is reasonable for us to limit our discussion to the contraction phase because the deceleration happens during the contraction phase only. Historically it has also been shown that the relaxation of *Spirostomum ambiguum* is an active process that requires ATP(78), and it is probably out of the scope of our model. Practically, our dynamically similar model also requires a small time step and makes simulating the slow relaxation phase prohibitively costly (see SI Appendix, Section A.7.5). For these reasons, we did not simulate the relaxation process. Finally, extending our model to 3D would require a new definition of the degree of entanglement. In 2D particle systems, entangled topology can be achieved by strings with point attachments to the particles, and we can quantify the degree of entanglement by the filament fraction defined in this manuscript. In 3D, however, a definition based on the reduced total degree of freedom is likely required, as two particles can now be connected by strings or sheets, which can give different degrees of constraint. Despite these differences, we expect our observations in 2D that increased entanglement in the system can create a strain-induced jamming material, help dampen the kinematics, and enhance the mechanical strength of the system can be readily extended to 3D cases. Also, our table-top experiments of table tennis balls and fabrics are a 3D realization of the concept, demonstrating the concept is transferable across 2D and 3D.

## Conclusions

In conclusion, we have discovered a unique entangled ER-vacuolar meshwork topology in *Spirostomum ambiguum*, with a fenestrated ER sheet spanning the entire millimeter-scale single-celled organism, wrapping around the vacuoles. Our work provides a new perspective on the potential role of ER geometry as a metamaterial critical for the survival of a cell undergoing rapid acceleration/deceleration cycles. Through modeling and table-top experiments, we demonstrate that the topological constraints from the entanglement can provide a braking/damping function, increase squeeze flow viscosity and help preserve internal spatial relationship against extreme motility, by strain-induced jamming transition. Using velocity direction spatial correlation of vacuolar meshwork, we confirm the presence of intracellular strain-induced jamming transition in *Spirostomum ambiguum* during its ultrafast contraction. Our study suggests a new mechanical role of RER, demonstrates a framework for gigantic cells to organize their organelle architecture efficiently and according to their evolutionary need, and points out a general design principle for a new class of shock-absorbing materials.

## Materials and Methods

### Protocols to culture *Spirostomum ambiguum*

*Spirostomum ambiguum* was cultured in spring water infused with Timothy hay and boiled wheat grains(79). See SI Appendix, Section A.1.

### Live cell imaging and high speed imaging

The organisms were introduced in a rectangular microfluidic chip of glass slides with copper side walls which act as electrodes. A Phantom V1210 high-speed camera mounted to a Nikon ECLIPSE Ti2 microscope and an electrophysiological DC pulse generator were synchronized with a micro-controller. See SI Appendix, Section A.2 & A.3.

### Transmission electron microscopy

The TEM sections of relaxed and contracted organisms were either imaged using a Tecnai 12 120kV TEM and data recorded using an UltraScan 1000 with Digital Micrograph 3 software at UC Berkeley, or imaged in the JEOL JEM-1400 120kV with photo taken using a Gatan Orius 2k X 2k digital camera at Stanford University. See SI Appendix, Section A.4 & A.5 for detailed sample preparations.

### Confocal microscopy

The confocal imaging was done on an inverted Zeiss LSM 780 multiphoton laser scanning confocal microscope. See SI Appendix, Section A.6 for detailed sample preparations.

### Detailed model description

See SI Appendix, Section A.7 for the detailed model descriptions. This includes the model overview (A.7.1), basic definitions (A.7.2), description of energy terms (A.7.3), determination of parameters (A.7.4), boundary movement and time step selections (A.7.5), practical considerations of running the simulations (A.7.6), details on tracing neighboring relationship (A.7.7), calculation of squeeze flow viscosity (A.7.8), validations of the models (A.7.9), the biological relevance of the model (A.7.10), and sensitivity testing to simulation parameters (A.7.11).

### Table-top experiments

We used table tennis balls wrapped by sheer fabric backdrop curtains. In the topologically constrained group, the fabric was cut into interconnected stripes. 45 table tennis balls were randomly attached to stripes in an interwoven fashion. In the control group, the fabric was joined to form a bag, and 45 table tennis balls were put into the bag. See SI Appendix, Section A.8.

## Data, Materials, and Software Availability

The data that support the plots within this paper and other findings of this study are available on BioImage Archive (access number: S-BIAD728). The code used for simulation and analysis is available on Github (jrchang612/ Entangled Soft Particles).

## ACKNOWLEDGMENTS

We thank all members of the Prakash Lab for scientific discussions and comments, including A. J. T. M. Mathijssen, H. Li, D. Krishnamurthy, E. Flaum, and S. Banvar. We thank R. Konte, R. Chajwa, and J. Zhu for the figure graphics and video. We thank Dr. D. Jorgens and R. Zalpuri at the University of California Berkeley Electron Microscope facility service for advice and assistance in electron microscopy and staff of Stanford Cell Sciences Imaging Facility (ARRA Award Number 1S10RR026780-01), Stanford Nano Shared Facilities (NSF award ECCS-2026822). We thank Professor C. S. O’Hern and J. Treado for the helpful discussion on modeling particle systems. We thank D. Krishnamurthy for providing his object-oriented code base, and we thank Professor P. Nuyujukian for providing the singularity container. RC gratefully acknowledges the support of Stanford University Bio-X SIGF Fellows Program and by the Ministry of Education in Taiwan. This work was supported by HHMI Faculty Fellows Award (M.P), BioHub Investigator Fellowship (M.P), Keck Foundation Research Grant, and NSF CCC (DBI1548297).

## Notes

The authors report no conflict of interest.

https://www.ebi.ac.uk/biostudies/bioimages/studies/S-BIAD728?query=S-BIAD728

https://github.com/jrchang612/Entangled_Soft_Particles

### Competing Interest Statement

The authors have declared no competing interest.

### Summary of Updates

1. the entangled hard particle model is now updated to entangled soft particle model, and we used dimensionless matching to ensure dynamic similarity of our model. 2. We used spatial correlation of velocity direction to demonstrate strain-induced jamming transition inside Spirostomum ambiguum.

## References

1. AM Wilson, et al., Locomotion dynamics of hunting in wild cheetahs. Nature 498, 185–189 (2013).

2. NA Yaraghi, et al., The Stomatopod Telson: Convergent Evolution in the Development of a Biological Shield. Adv. Funct. Mater. 29, 1902238 (2019).

3. NS Ha, G Lu, DW Shu, TX Yu, Mechanical properties and energy absorption characteristics of tropical fruit durian (Durio zibethinus). J. Mech. Behav. Biomed. Mater. 104, 103603 (2020).

4. MA Ferenczi, et al., Why muscle is an efficient shock absorber. PLoS ONE 9, e85739 (2014).

5. TT Li, H Wang, SY Huang, CW Lou, JH Lin, Bioinspired foam composites resembling pomelo peel: Structural design and compressive, bursting and cushioning properties. Compos. Part B: Eng. 172, 290–298 (2019).

6. F Stein, Der Organismus der Infusionsthiere. (W. Engelmann,, Leipzig,), pp. 187–207 (1867).

7. AJ Mathijssen, J Culver, MS Bhamla, M Prakash, Collective intercellular communication through ultra-fast hydrodynamic trigger waves. Nature 571, 560–564 (2019).

8. T Nüchter, M Benoit, U Engel, S Özbek, Holstein, Nanosecond-scale kinetics of nematocyst discharge. Curr. Biol. 16, R316–R318 (2006).

9. H Ishida, Y Shigenaka, Cell model contraction in the ciliate spirostomum. Cell Motil. Cytoskelet. 9, 278–282 (1988).

10. RB Hawkes, DV Holberton, Myonemal contraction of spirostomum. II. Some mechanical properties of the contractile apparatus. J. Cell. Physiol. 85, 595–602 (1975).

11. A Upadhyaya, MP Sheetz, Tension in Tubulovesicular Networks of Golgi and Endoplasmic Reticulum Membranes. Biophys. J. 86, 2923–2928 (2004).

12. S Gogia, S Neelamegham, Role of fluid shear stress in regulating VWF structure, function and related blood disorders in Biorheology. (IOS Press) Vol. 52, pp. 319–335 (2015).

13. RHJ Brown, The Protoplasmic Viscosity of Paramecium. J. Exp. Biol. 17, 317–324 (1940).

14. SS Rogers, TA Waigh, JR Lu, Intracellular Microrheology of Motile Amoeba proteus. Biophys. J. 94, 3322 (2008).

15. H Ishida, T Suzaki, Y Shigenaka, Cell body contraction of Spirostomum does not involve shortening of inter-kinetosomal distance along ciliary lines. Zool. Sci. 13, 669–672 (1996).

16. A Bishop, Memoirs: Some Observations Upon Spirostomum ambiguum (Ehrenberg). J. Cell Sci. s2-67, 391–434 (1923).

17. W Foissner, H Berger, A user-friendly guide to the ciliates (Protozoa, Ciliophora) commonly used by hydrobiologists as bioindicators in rivers, lakes, and waste waters, with notes on their ecology, Technical Report 7996 (1991).

18. JA Raven, The Vacuole: a Cost-Benefit Analysis. Adv. Bot. Res. 25, 59–86 (1997).

19. C Laporte, et al., A necrotic cell death model in a protist. Cell Death & Differ. 14, 266–274 (2006).

20. A Shubin, I Demidyuk, A Komissarov, L Rafieva, S Kostrov, Cytoplasmic vacuolization in cell death and survival. Oncotarget 7, 55863–55889 (2016).

21. T Yamanaka, N Nukina, ER Dynamics and Derangement in Neurological Diseases. Front. Neurosci. 12, 91 (2018).

22. L Scorrano, et al., Coming together to define membrane contact sites. Nat. Commun. 10, 1–11 (2019).

23. KM Browne, X. —The Golgi apparatus and other cytoplasmic bodies in Spirostomum ambiguum. J. Royal Microsc. Soc. 58, 188–199 (1938).

24. JES Moore, On the Structural Differentiation of the Protozoa as seen in Microscopic Sections. Zool. J. Linnean Soc. 24, 364–368 (1893).

25. P Ramoino, A Diaspro, M Fato, F Beltrame, M Robello, Changes in the endoplasmic reticulum structure of Paramecium primaurelia in relation to different cellular physiological states. J. Photochem. Photobiol. B: Biol. 54, 35–42 (2000).

26. S Phonekeo, N Mlot, D Monaenkova, DL Hu, C Tovey, Fire ants perpetually rebuild sinking towers. Royal Soc. Open Sci. 4 (2017).

27. JL Shivers, et al., Compression stiffening of fibrous networks with stiff inclusions. Proc. Natl. Acad. Sci. 117, 21037–21044 (2020).

28. N Gravish, SV Franklin, DL Hu, D. Goldman, Entangled Granular Media. Phys. Rev. Lett. 108, 208001 (2012).

29. E Panagiotou, KC Millett, PJ Atzberger, Topological Methods for Polymeric Materials: Characterizing the Relationship Between Polymer Entanglement and Viscoelasticity. Polym. 2019, Vol. 11, Page 437 11, 437 (2019).

30. D Parisi, et al., Nonlinear Shear Rheology of Entangled Polymer Rings. Macromolecules 54, 2811–2827 (2021).

31. P Aejmelaeus-Lindström, J Willmann, S Tibbits, F Gramazio, M Kohler, Jammed architectural structures: Towards large-scale reversible construction. Granul. Matter 18, 1–12 (2016).

32. BC White, A Garland, R Alberdi, BL Boyce, Interpenetrating lattices with enhanced mechanical functionality. Addit. Manuf. 38, 101741 (2021).

33. A Boromand, A Signoriello, F Ye, C. O’Hern, MD Shattuck, Jamming of Deformable Polygons. Phys. Rev. Lett. 121, 248003 (2018).

34. J Zhang, TS Majmudar, M Sperl, RP Behringer, Jamming for a 2D granular material. Soft Matter 6, 2982–2991 (2010).

35. S Shankar, L Mahadevan, Active muscular hydraulics. bioRxiv p. 2022.02.20.481216 (2022).

36. R Dimova, CM Marques, eds., The giant vesicle book. (CRC Press, Taylor & Francis Group), pp. 85, 284–302 (2019).

37. EM Ettienne, Control of contractility in Spirostomum by dissociated calcium ions. J. Gen. Physiol. 56, 168–179 (1970).

38. Y Wu, et al., Contacts between the endoplasmic reticulum and other membranes in neurons. Proc. Natl. Acad. Sci. 114, E4859–E4867 (2017).

39. M Van Hecke, Jamming of soft particles: geometry, mechanics, scaling and isostaticity. J. Physics: Condens. Matter 22, 033101 (2009).

40. L Meijering, Interface area, edge length, and number of vertices in crystal aggregates with random nucleation. Philips Res. Reports 8, 270–290 (1953).

41. Y Fujii, et al., Spontaneous spatial correlation of elastic modulus in jammed epithelial monolayers observed by AFM. Biophys. J. 116, 1152–1158 (2019).

42. HE Finley, Electron microscopy of thin-sectioned spirostomum. Science 113, 362–363 (1951).

43. HE Finley, Electron microscopical observations on Spirostomum ambiguum. Annals New York Acad. Sci. 62, 231–246 (1955).

44. R Yagiu, Y Shigenaka, Electron Microscopy of the Longitudinal Fibrillar Bundle and the Contractile Fibrillar System in Spirostomum ambiguum. The J. Protozool. 10, 364–369 (1963).

45. HE Finley, CA Brown, WA Daniel, Electron Microscopy of the Ectoplasm and Infraciliature of Spirostomum ambiguum. The J. Protozool. 11, 264–280 (1964).

46. WA Daniel, CF Mattern, Some Observations on the Structure of the Peristomial Membranelle of Spirostomum ambiguum. The J. Protozool. 12, 14–27 (1965).

47. WJ Lehman, LI Rebhun, The structural elements responsible for contraction in the ciliate Spirostomum. Protoplasma 72, 153–178 (1971).

48. VS Hobbs, RA Jenkins, JR Bamburg, Evidence for the lack of actin involvement in mitosis and in the contractile process in Spirostomum teres. J. Cell Sci. 60, 169–179 (1983).

49. Y Shigenaka, J Hosoi, M Ando, M Ishida, I Ishii, Ultrastructure of the Isolated Microtubules and Intermicrotubular Bridges in a Heterotrichous Ciliate, Spirostomum ambiguum. J. Electron Microsc. 38, 363–370 (1989).

50. FG Pautard, Hydroxyapatite as a developmental feature of Spirostomum ambiguum. Biochimica et Biophys. Acta 35, 33–46 (1959).

51. PR Burton, Fine Structure of Mitochondria of Spirostomum ambiguum as Seen in Sectioned and Negatively-Stained Preparations. The J. Protozool. 17, 295–299 (1970).

52. DN Harrison, CH Dorsey, CA Brown, Studies on a macronuclear endosymbiont of Spirostomum ambiguum. II. Ultrastructural comparison of the in situ and the cultivated endosymbiont. Transactions Am. Microsc. Soc. 95, 565–568 (1976).

53. SI Fokin, M Schweikert, F Brümmer, HD Görtz, Spirostomum spp. (Ciliophora, Protista), a suitable system for endocytobiosis research. Protoplasma 225, 93–102 (2005).

54. V Fallon, PE Garner, JE Aaron, Mineral Fabrication and Golgi Apparatus Activity in Spirostomum ambiguum: A Primordial Paradigm of the Stressed Bone Cell? J. Biomed. Sci. Eng. 10, 466–483 (2017).

55. D Osborn, TC Hamilton, Electron microbeam analysis of calcium distribution in the ciliated protozoan, Spirostomum ambiguum. J. Cell. Physiol. 91, 409–416 (1977).

56. A Borsellino, B Cavazza, M Riani, Photodynamic action in Spirostomum ambiguum. Biophysik 8, 30–41 (1971).

57. A Le Dû-Delepierre, G Persoone, C. Groliére, A new low cost microbiotest with the freshwater ciliate protozoan Spirostomum ambiguum: Definition of culturing conditions. Hydrobiologia 325, 121–130 (1996).

58. J Pérez-Sancho, et al., The arabidopsis synaptotagmin1 is enriched in endoplasmic reticulum-plasma membrane contact sites and confers cellular resistance to mechanical stresses. Plant Physiol. 168, 132–143 (2015).

59. M Araújo, A Tavares, DV Vieira, IA Telley, RA Oliveira, Endoplasmic reticulum membranes are continuously required to maintain mitotic spindle size and forces. Life Sci. Alliance 6 (2023).

60. H Merta, et al., Cell cycle regulation of ER membrane biogenesis protects against chromosome missegregation. Dev. cell (2021).

61. I Mukhtar, et al., Transcriptome Profiling Revealed Multiple rquA Genes in the Species of Spirostomum (Protozoa: Ciliophora: Heterotrichea). Front. Microbiol. 11, 3373 (2021).

62. TD Mattia, et al., Identification of MOSPD2, a novel scaffold for endoplasmic reticulum membrane contact sites. EMBO reports 19, e45453 (2018).

63. K Bradonjić, D Swain, A Widom, N Srivastava, The Casimir Effect in Biology: The Role of Molecular Quantum Electrodynamics in Linear Aggregations of Red Blood Cells. J. Physics: Conf. Ser. 161, 012035 (2009).

64. L Wu, LY Lanry Yung, KM Lim, Dielectrophoretic capture voltage spectrum for measurement of dielectric properties and separation of cancer cells. Biomicrofluidics 6, 014113 (2012).

65. LH Klausen, T Fuhs, M Dong, Mapping surface charge density of lipid bilayers by quantitative surface conductivity microscopy. Nat. Commun. 7, 1–10 (2016).

66. H Hao, et al., The nesprin-1/-2 ortholog ANC-1 regulates organelle positioning in c. elegans independently from its KASH or actin-binding domains. Elife 10 (2021).

67. H Ishida, T Suzaki, C Kuribayashi, E Masuyama, O Numata, Distribution of Actin-like Proteins in the Ciliate Spirostomum ambiguum. Jpn. J. Protozool. 36, 141–146 (2003).

68. J Zhang, et al., Giant proteins in a giant cell: Molecular basis of ultrafast Ca2+-dependent cell contraction. Sci. Adv. 9 (2023).

69. AJ Liu, SR Nagel, Jamming is not just cool any more. Nature 396, 21–22 (1998).

70. D Bi, J Zhang, B Chakraborty, RP Behringer, Jamming by shear. Nature 480, 355–358 (2011).

71. L Zorzetto, D Ruffoni, Wood-Inspired 3D-Printed Helical Composites with Tunable and Enhanced Mechanical Performance. Adv. Funct. Mater. 29, 1805888 (2019).

72. M Zou, S Xu, C Wei, H Wang, Z Liu, A bionic method for the crashworthiness design of thin-walled structures inspired by bamboo. Thin-Walled Struct. 101, 222–230 (2016).

73. D Ruan, G Lu, B Wang, TX Yu, In-plane dynamic crushing of honeycombs - A finite element study. Int. J. Impact Eng. 28, 161–182 (2003).

74. DK Rajak, L Kumaraswamidhas, S Das, An Energy Absorption Behaviour of Foam Filled Structures. Procedia Mater. Sci. 5, 164–172 (2014).

75. J Rivera, et al., Toughening mechanisms of the elytra of the diabolical ironclad beetle. Nature 586, 543–548 (2020).

76. J Zhang, G Lu, Z You, Large deformation and energy absorption of additively manufactured auxetic materials and structures: A review. Compos. Part B: Eng. 201, 108340 (2020).

77. TL Jahn, Contraction of protoplasm. II. Theory: Anodal vs. Cathodal in relation to calcium. J. Cell. Physiol. 68, 135–148 (1966).

78. H Ishida, T Suzaki, Y Shigenaka, Effect of Mg2+ on Ca2+-dependent contraction of a Spirostomum cell model. Eur. J. Protistol. 32, 316–319 (1996).

79. PJ Hummer, Culturing & Using Protozoans in the Laboratory. Am. Biol. Teach. 55, 357–360 (1993).

