## Supplementary material for "Topological damping in an ultrafast giant cell": SI Appendix

#### This PDF file includes:

Supporting text
Figs. S1 to S12
Table S1
Legends for Movies S1 to S13
SI References

#### Other supporting materials for this manuscript include the following:

Movies S1 to S13

### Supporting Information Text

**B Perturbation experiments using ER drugs** **7**

**C Further discussions on volume fractions of vacuoles** **8**

**D Discussions on lipid droplets** **8**

**E The discrepancy of vacuole deformations between simulations and experiments** **8**

### A. Materials and methods

**A.1. Culture.** *Spirostomum ambiguum* (Sciento catalog #P346 and Ward's Science catalog #470176-728) was cultured in Hummer medium (1) of spring water infused with Timothy hay (Carolina catalog #132385) and boiled wheat grains (Carolina catalog #132425) as previously described(2). Organisms used in experiments were extracted in the exponential growth phase. The organisms from Sciento (the company ceased trading in 2021) were used for the transmission electron microscopy, and the organisms from Ward's Science were used for the confocal microscopy.

**A.2. Live cell imaging and high speed imaging.** The live imaging was performed as previously described method(2). In brief, the organisms were introduced in a rectangular microfluidic chip (length 2.2cm, width 1cm, height 150$\mu$m) of glass slides with copper side walls which act as electrodes. A high speed camera (Phantom V1210 camera, 10000fps) mounted to the microscope (Nikon ECLIPSE Ti2, 40X DIC objective) and an electrophysiological DC pulse generator (Grass instruments, S88) were synchronized with a micro-controller (Arduino Uno). Pulse voltage and duration were chosen according to previously reported results(2).

The live imaging of gelatin-embedded cells was performed as previously described method(3). In brief, *Spirostomum ambiguum* was introduced to 2% gelatin (G-Biosciences catalog #RC-053) solution in 1X Carter's media maintained at 30°C in a water bath. After thorough mixing, the gelatin solution containing the organism was introduced to the imaging chamber (glass slides and coverslips with spacer height 150 $\mu$m), and the imaging chamber was put on ice with glass slides facing down for 1-2 minutes. Monitor the cell and cilia activities under dissection scope while the imaging chamber is on ice. Remove the slides from the ice bath when the cells are no longer motile yet retain cilia activities.

**A.3. Estimation of vacuole strain from live cell imaging.** As the contraction of *Spirostomum ambiguum* has a torsional component, which brings vacuoles away from the focal plane, it is very difficult to track enough number of vacuoles before and after each contraction. Also due to the extremely high speed nature of the contraction process, it is hard to perform 3D imaging. We thus estimated the strain of the vacuoles based on the population statistics of the aspect ratio (AR), and not necessarily keeping track of the same vacuoles. Assuming that the deformation of vacuoles originates from a uniaxial compression force, and assuming that the volume of vacuoles is conserved during the contraction process, we can write the strain of the vacuoles in the direction parallel and perpendicular to the compressing force as $\varepsilon_3$ and $\varepsilon_1$, respectively. Based on the assumptions, we can also write $\varepsilon_3 < 0$, $\varepsilon_1 > 0$, $\varepsilon_3 + 2\varepsilon_1 = 0$, $\mathrm{AR} = (1 - 1/2\varepsilon_3)/(1 + \varepsilon_3)$. We thus can estimate the magnitude of compression strain by measuring the aspect ratio of the vacuoles before and after the contraction and calculating it with $\varepsilon_3 = (1 - \mathrm{AR})/(1/2 + \mathrm{AR})$.

**A.4. Fixing *Spirostomum ambiguum* in relaxed state.** On a glass coverslip, mix *Spirostomum ambiguum* with ice-cold 2X PHEM buffer in 1:1 proportion and put in 4°C immediately for 15 minutes. Gently mix *Spirostomum ambiguum* in 1X PHEM buffer with ice-cold 4% glutaraldehyde, 8% paraformaldehyde in 1X PHEM buffer again in 1:1 proportion, and put in 4°C immediately for at least 1 hour. Directly fixing the cells inside an eppendorf or on a 96-well plate is not recommended, as fixed organisms tend to adhere to the surface and will make the retrieval of fixed samples difficult.

**A.5. Transmission Electron Microscopy.** *Spirostomum ambiguum* were fixed in 2% glutaraldehyde with 4% paraformaldehyde in 1X PHEM buffer(4) for at least one hour at room temperature. The samples were stained with Safranin-O and stabilized in 1% low melting point agarose, then cut into 1mm cubes. Samples were rinsed (3×; 10 min, room temperature (RT)) in 1X PBS, and immersed in 1% osmium tetroxide with 1.6% potassium ferricyanide in 1xPBS, pH 7.4 for 1 hour. Samples were rinsed (3×; 10 min, RT) in buffer and then briefly in distilled water. Samples were then subjected to an ascending acetone gradient (10min; 35%, 50%, 70%, 80%, 90%) followed by pure acetone (2×; 10 min, RT). Samples were progressively infiltrated with Epon resin (EMS, Hatfield, PA, USA), while rocking, and then polymerized at 60℃ for 24-48 hours in silicon molds. Thick sections (200 nm) were cut using a Leica UC6 ultramicrotome (Leica, Wetzlar, Germany), collected on glass slides and used for general orientation. Thin sections (70-80 nm) were collected onto formvar-coated 50 mesh copper grids. Thin sections were post-stained with 2% aqueous uranyl acetate and Reynold's lead citrate, 5 mins each. Then sections were either imaged using a Tecnai 12 120kV TEM (FEI, Hillsboro, OR, USA) and data recorded using an UltraScan 1000 with Digital Micrograph 3 software (Gatan Inc., Pleasanton, CA, USA) at UC Berkeley, or imaged in the JEOL JEM-1400 120kV with photo taken using a Gatan Orius 2k X 2k digital camera at Stanford University. The montage image was created by SerialEM. All segmentation of organelles was done manually on Fiji.

**A.6. Confocal Imaging.** *Spirostomum ambiguum* were stained alive in 625 ng/mL DiOC6(3) (Sigma-Aldrich catalog #318426-250MG) and 500 nM MitoTracker™ Orange CMTMRos (Thermo Fisher Scientific catalog #M7510) for 30 minutes, followed by washing steps in 1X Carter's media(5). The cells were then fixed in Bouin's solution for 30 minutes, and washed in 1X Carter's media until the supernatant became completely transparent and without any yellow-tinged color. The fixed samples are then stained with Hoechst 33342 (Invitrogen catalog #H3579) in 1:1000 dilution for 15 minutes, followed by washing steps in 1X Carter's media. The fixed samples were transferred to a glass slide, mounted in Vectashield antifade mounting media (Vector labs catalog #H-1000), and sealed with #1.5 coverslip. All Chemicals were diluted from stock solution using 1X Carter's media.

For tunicamycin experiments, *Spirostomum ambiguum* were treated in 1 $\mu$g/mL tunicamycin (Millipore Sigma catalog #T7765, stock solution in DMSO, 1:1000 dilution) in 1X Carter's media overnight (higher concentration such as 5 $\mu$g/mL kills the organisms). The cells were stained alive in 625 ng/mL DiOC6(3) (Sigma-Aldrich catalog #318426-250MG) and 500 nM MitoSpy™ Orange CMTMRos (BioLegend catalog #424803) for 30 minutes, followed by washing steps in 1X Carter's media. The cells were then fixed in Bouin's solution for 30 minutes, and washed in 1X Carter's media until the supernatant became completely transparent and without any yellow-tinged color. The fixed samples were then stained with Hoechst 33342 (Invitrogen catalog #H3579) in 1:1000 dilution for 15 minutes, followed by washing steps in 1X Carter's media. The fixed samples are transferred to a glass slide, mounted in Vectashield antifade mounting media (Vector labs catalog #H-1000), and sealed with #1.5 coverslip. All Chemicals were diluted from stock solution using 1X Carter's media.

For Nile red experiments, *Spirostomum ambiguum* were stained alive in 625 ng/mL DiOC6(3) for 30 minutes, followed by washing steps in 1X Carter's media. The cells were then fixed in Bouin's solution for 30 minutes, and washed in 1X Carter's media until the supernatant became completely transparent and without any yellow-tinged color. The fixed samples are then stained with 6$\mu$M Nile red (TCI Chemicals catalog #N0659, stock solution in DMSO, 1:1000 dilution) and Hoechst 33342 in 1:1000 dilution for 15 minutes, followed by washing steps in 1X Carter's media. The fixed samples are transferred to a glass slide, mounted in Vectashield antifade mounting media (Vector labs catalog #H-1000), and sealed with #1.5 coverslip. All Chemicals are diluted from stock solution using 1X Carter's media.

We used Bouin's solution as our fixatives because of its better preservation of soft and delicate structures(6). However, as the picric acid is a strong oxidizer and may impact the fluorophores, we used a much shorter fixation time, and all the imaging was done within 24 hours after the fixation. The confocal imaging was done on an inverted Zeiss LSM 780 multiphoton laser scanning confocal microscope, using PLAN APO 63X oil objective (NA 1.4).

**A.7. Model description.**

***A.7.1. Model overview.*** In the entangled soft particle model, the cell is simplified into a two-dimensional rectangle, the vacuoles in the cytoplasm are simplified into soft particles, and the RER is simplified into flexible strings, with resistance to bending and stretching, and a critical stretching strain before breaking. The total number of soft particles and strings are $N_\mathrm{c}$ and $N_\mathrm{f}$, respectively. The soft particles are simulated following the vertex-based deformable particle model with the smooth surface method as explained by Boromand et al.(7) and Vanderwerf et al.(8), with adjustment to include entangled strings, their associated energy, and repulsive energy from the boundaries. (Note that the rough surface method, the other simulation method used in Boromand et al., is not appropriate as it would introduce friction among particles when there is bidispersity.) Each particle is discretized into polygons with $N_\mathrm{v}$ vertices. Each string is discretized into $N_\mathrm{p}$ points (and thus $(N_\mathrm{p} - 1)$

segments), and the natural length of each segment is $dl_{\mathrm{f}}$. The total length of the string is thus $l_{\mathrm{f}} = (N_{\mathrm{p}} - 1)dl_{\mathrm{f}}$. The first and last discretized points of each string are attached to the surface of the two particles and are fixed relative to the attached particles. There is no energy penalty associated with the changes in the exit angle between the string segment and the surface of the particle. The forces exerted on the first or last segment of each string will be felt by the attached particles, which can cause translation and rotation. In the following, we will first describe the energy terms involved in the simulation, followed by the description of how we determine all parameters by matching the relevant dimensionless numbers, and we conclude by discussing the practical tips we used that can help the simulation finish in a reasonable amount of time. (Fig. S3A&B)

***A.7.2. Basic definitions of the model.*** In our system, the position of the $i$-th vertex of $m$-th particle is $\vec{x}^v_{m,i}$, and the vector of the $i$-th edge of the $m$-th particle is $\vec{l^c_{m,i}} := \vec{x}^v_{m,i+1} - \vec{x}^v_{m,i}$, with length $l^c_{m,i}$. The radius of the $m$-th particle is $r_m$, and the sizes of the soft particles have a bidispersity of 1.4, with an exact 50%/50% ratio(9). Therefore, the radius of each particle $m$ can either be $R_{\mathrm{c}}$ or $1.4R_{\mathrm{c}}$, with $R_{\mathrm{c}}$ as the baseline radius defined for scaling of other length scales in the system. The edges of the soft particles are simulated as circulo-lines with width $\delta = 2R_{\mathrm{c}}\sin(\pi/N_{\mathrm{v}})$, and we will detail the formula to calculate repulsive energy shortly. (Fig. S3A&B)

We chose 1:1.4 bidisperse deformable particle because it has the lowest critical volume fraction for jamming transition, and it is also the most widely studied 2D particle mixture in the literature of jamming transition(9). This allows us to establish entangled topology as a general strategy to create strain-induced jamming transitions, rather than just trying to fit a specific type of particle distribution. This also allows us to compare our results with the literature. The experimental size distribution of vacuoles is shown in Figure S9. It is polydisperse, but we don't expect the conclusions to change if we use an experimentally calibrated size distribution. The critical volume fraction for the jamming transition will be different, but the topologically-assisted strain-induced jamming transition described in the main text should be the same.

The cell in its neutral shape is a rectangular box with side length $L_x$ and $L_y$. The volume fraction of the system (the space occupied by the circulo-lines are included) $\phi$ can be easily computed as $\phi = N_{\mathrm{c}}(\frac{N_{\mathrm{v}}}{2}\sin(\frac{2\pi}{N_{\mathrm{v}}})R_{\mathrm{c}}^2(\frac{1^2+1.4^2}{2}) + \frac{1}{2}\delta^2 N_{\mathrm{v}}(\frac{1+1.4}{2}) + \pi(\frac{\delta}{2})^2)/(L_x L_y)$. The natural length of each string is $1.4 \times \mathrm{mLf} \times R_{\mathrm{c}}$. The prefactor mLf was chosen to be 4.8 for the simulations reported in the paper. We have also run the simulation with $\mathrm{mLf} = 3.2$ or 6.4, and the conclusions are similar. These numbers are chosen to be on the same order as $\pi$, such that each string can wrap around 2 large particles without extension, with the attached positions on the opposite side (e.g., one end at the 3 o'clock position of the first particle and the other end at the 9 o'clock position of the second particle) to mimic the topology observed under TEM. The natural length of each string is a tune-able parameter, but the inequality $l_{\mathrm{f}} > \sqrt{\frac{L_x L_y}{N_{\mathrm{c}}}}$ must hold to ensure that the particles are not constricted into a smaller subspace of the entire cell. The position of the $i$-th discretization point of the $n$-th string is $\vec{x}_{n,i}$, and the length of the $i$-th segment of the discretized string is $l_{n,i} := |\vec{x}_{n,i+1} - \vec{x}_{n,i}|$. The discretization points of the string and the vertices of the soft particle have no size, and this will require adjustment on the repulsion energy terms compared to Boromand et al(7). We eliminated the size of the discretization points of the string so that when we increase the number of strings in the system, the volume fraction will not change. By doing so, we can ensure all the changes are caused by the topological constraints alone.

***A.7.3. Relevant energy terms.*** The system evolves according to an over-damped molecular dynamic scheme(10), and the energy of the system $U$ is composed of 10 components (Fig. 3A, Fig. S3C), which can be divided into three categories: (1) repulsive energy terms, which include particle-particle repulsive energy ($U_{\mathrm{r}}$), particle-string repulsive energy ($U_{\mathrm{rl}}$), particle-boundary repulsive energy ($U_{\mathrm{rb}}$), and string-boundary repulsive energy ($U_{\mathrm{rbf}}$). (2) single-particle energy terms, which include particle compressibility term ($U_{\mathrm{a}}$), particle surface tension term ($U_{\gamma}$), particle surface contractility term ($U_{\mathrm{lc}}$), and the particle surface bending term ($U_{\mathrm{b}}$). (3) single-string energy terms, which include string contractility term ($U_{\mathrm{l}}$) and string bending term ($U_{\mathrm{bf}}$). Detailed definitions of the functions related to the geometry between points and segments can be found in Figure S3B. The particles have no Brownian motion, as we have never observed significant Brownian motion of the vacuoles in the cytoplasm. Note that we did not include the repulsive energy of strings crossing over each other because this is a 2D approximation of the 3D RER wrapping. The separate strings in 2D actually represent one large contiguous ER membrane in 3D, so they can cross over each other with zero energy penalty. Nonetheless, the same reasoning does not apply to particles, as separate particles in 2D represent separate vacuoles in 3D and can never cross over each other without energy penalty.

The gradient of the potential energy generates forces $\vec{F}$ on the vertices of particles and the discretization points of the strings. With $\eta$ as the viscosity, the velocity of the $i$-th vertex of $m$-th particle is computed as $\vec{v}^v_{m,i} = \sum \vec{F}^v_{m,i}/(\pi R_{\mathrm{c}}\eta/N_{\mathrm{v}})$. The velocity of the $i$-th discretization point of the $n$-th string is $\vec{v}_{n,i} = \sum \vec{F}_{n,i}/(\eta dl_{\mathrm{f}})$. We performed validation simulations on the shape relaxation of a single particle and filament to confirm that the solution from this formula has the expected time scale.

When the string is extended beyond the critical strain (see Section A.7.4 for the determination of parameters), the string ruptures, and the string contractility energy term of the ruptured string right before its rupture is added to the total sacrificial bond effect.

***A.7.4. Determination of parameters.*** The simulation has 13 parameters in total: initial boundary length in $x$ and $y$ directions ($L_x^{\mathrm{sim}}$, $L_y^{\mathrm{sim}}$), baseline radius of the vacuole ($r_{\mathrm{vac}}^{\mathrm{sim}}$, same as $R_{\mathrm{c}}$), characteristic velocity of the particle ($u^{\mathrm{sim}}$), viscosity ($\eta^{\mathrm{sim}}$), boundary force ($F^{\mathrm{sim}}$), duration of the contraction ($\tau^{\mathrm{sim}}$), length contractility constant ($k_{\mathrm{l}}^{\mathrm{sim}}$), area compressibility constant ($k_{\mathrm{a}}^{\mathrm{sim}}$), interfacial tension ($\gamma^{\mathrm{sim}}$), surface bending constant ($k_{\mathrm{b}}^{\mathrm{sim}}$), repulsivity constant ($k_{\mathrm{r}}^{\mathrm{sim}}$), and the critical extension before the rupture of the string ($\varepsilon_{\mathrm{crit}}^{\mathrm{sim}}$). To determine these parameters, we used 8 physical parameters related to the contraction kinematics, geometry of the organism, and fluid properties (length of the cell $L \sim 1$ mm, diameter of the cell $D \sim 100$ $\mu$m, radius of the vacuole $r_{\text{vac}} \sim 10\ \mu$m, boundary force $F \sim 10^{-6}$ N(11), characteristic velocity of the contraction $u \sim 0.2$ m/s(2), characteristic duration of the contraction $\Delta t \sim 5$ msec(2), viscosity of the cytoplasm $\eta_{\text{cyto}} \sim 0.005$ Pa-sec)(12) (measurement in other protists), and the mass density of the cytoplasm ($\rho \sim 1000$ kg/m$^3$). We also performed a literature search to find 6 parameters related to the mechanical properties of the vacuoles (area expansion modulus $k_{\text{A}}$ ∼200 mN/m(13), water bulk modulus $B \sim 2.2 \times 10^9$ Pa(14), interfacial tension $\gamma \sim 50$ mN/m(13), bending modulus $\kappa \sim 10^{-19}$ J(13), hydration repulsion pressure of lipid bilayer membrane $p_0 \sim 3.6 \times 10^{-4}$ Pa(15), and the critical stretching strain of lipid bilayer membrane before rupture $\varepsilon_{\text{crit}} \sim 3\%$(16)). We then compute 10 dimensionless numbers using the contraction kinematics and the membrane mechanical properties (Reynolds number based on vacuole radius $\text{Re}_r$, Stokes number based on external force $\text{Stk}_F$, Strouhal number St, capillary number based on membrane stretching Ca, aspect ratio of the cell AR, aspect ratio between the length of the cell and the radius of the vacuole $\text{AR}_{\text{vac}}$, and the four relative strength ratio between the 5 membrane mechanical properties $K_1$, $K_2$, $K_3$, and $K_4$, following the definitions in Boromand et al(7)). Among these dimensionless numbers, the extremely small number of $K_1$ introduced huge stiffness to the differential equation, and even implicit methods frequently fail to converge on the solution. We therefore relax that constraint by multiplying $K_1$ 100 times (we will explain later through time scale analysis and show that this adjustment still preserves the physics). After arbitrarily setting the radius of the vacuole and the characteristic velocity in the simulation (these 2 set the magnitude of simulation length and time scale), all the remaining parameters are determined by matching the 10 dimensionless numbers.

Note that as we previously described, there was no reported measurement of the cytoplasm viscosity in *Spirostomum*, and the previously measured cytoplasm viscosity in other ciliates or protists fall in the range of 0.005-0.05 Pa-sec(12, 17). We have also performed our simulation assuming a cytoplasm viscosity of 0.05 Pa-sec. However, in this scenario, even the system with 0 filament fraction ($\phi_{\text{f}}$) can only contract itself to 70-80% of its original body length within 5 msec (Fig. S8). We thus believe the actual cytoplasm viscosity of *Spirostomum* should be on the lower end of this range.

***A.7.5. Boundary movement and time step selections.*** As we are only interested in the damping during the contraction phase, we only perform simulation during the contraction phase and not the relaxation phase. In the contraction phase, a constant boundary force $F^{\text{sim}}$ is imposed, and the reactive forces of the system on the side walls $F_x$ are computed. The bounding box is then shrunken by $\Delta x = (F^{\text{sim}} - F_x)/(6\pi\eta L^{\text{sim}}_{y,0})$. Note that the $\eta$ we used here is the viscosity of water converted to simulation units using dimensionless number matching. The boundary evolution is incorporated into the differential equation, and the bounding box size in $y$-direction is adjusted accordingly to ensure area conservation. The simulation stops when it reaches the duration of the contraction ($\tau^{\text{sim}}$), or when the box is contracted to <5% of its original length in $x$-direction. At each time step, the net energy input is calculated by $(F^{\text{sim}} - F_x)\Delta x$, and its cumulative sum over time reports the cumulative net energy input. The code is written in Python3 and the systems of ODEs are solved deterministically by the backward differentiation formula (BDF) method of order 5 (18) (with atol = 1E-7 and rtol = 1E-6) of the package odespy(19).

To ensure accurate simulation results, we need to analyze the relevant time scale in the simulation process and chose our time step accordingly. As listed in Table S1, when we are using implicit methods, we want our time step to be greater than the smallest time scale (the area relaxation time scale) to gain benefit from using implicit methods, but still smaller than the second smallest time scale (the repulsion relaxation time scale) to resolve the dynamics accurately. We perform sub-sampling to the simulation result to reduce the amount of data that needs to be saved. Note that the length relaxation time scale and repulsion relaxation time scale are on the same order of magnitude, and that suggests there will be an interaction between particle surface stretching and particle repulsion, which requires a smaller time step to resolve. The final time step used for simulation is thus at least 20 times smaller than the second-smallest relaxation time scale (in this case, the repulsion relaxation time scale).

**Table S1. Time scale analysis of the simulation.**

| - | high viscosity | low viscosity |
|---|---|---|
| cytoplasm viscosity | 0.05 Pa-sec | 0.005 Pa-sec |
| $\eta^{\text{sim}}$ | 250 | 25 |
| length relaxation time scale | 5 | 0.5 |
| area relaxation time scale | 4.5E-3 | 4.5E-4 |
| bending relaxation time scale | 1E9 | 1E8 |
| tension relaxation time scale | 20 | 2 |
| repulsion relaxation time scale | 2.78 | 0.278 |
| appropriate time step | 0.1 | 0.01 |

length relaxation time scale: $\tau = \eta^{\text{sim}} r^{\text{sim}}_{\text{vac}} / k^{\text{sim}}_{\text{l}}$.
area relaxation time scale: $\tau = \eta^{\text{sim}} / r^{\text{sim}}_{\text{vac}} k^{\text{sim}}_{\text{a}}$.
bending relaxation time scale: $\tau = \eta^{\text{sim}} (r^{\text{sim}}_{\text{vac}})^5 / k^{\text{sim}}_{\text{b}}$.
tension relaxation time scale: $\tau = \eta^{\text{sim}} (r^{\text{sim}}_{\text{vac}})^2 / \gamma^{\text{sim}}$.
repulsion relaxation time scale: $\tau = \eta^{\text{sim}} r^{\text{sim}}_{\text{vac}} / k^{\text{sim}}_{\text{r}}$.

***A.7.6. Practical considerations of running the simulations.*** As soft particle simulation contains a huge amount of degree of freedom, several simulation tricks were used to help finish the simulation in a reasonable amount of time. During the initialization phase, we used hard particle simulation, instead of soft particle simulation, to make the initialization phase faster. To initialize the system, the center locations of the particles are randomly initialized within the box according to a standard algorithm based on polygon partitioning. After a brief initial relaxation period without strings attached, a Delaunay triangulation map is then generated based on the center location of the particles to find the neighbors(20), with the total number of edges in the Delaunay triangulation map $N_{\mathrm{D}}$. The strings are initialized by randomly connecting the neighbors in the Delaunay triangulation map at the opposite positions, without replacement, so the same two particles will not be connected by more than one string. The phase angle of its attachment on one end, $\theta_1$, is randomly initialized between 0 and $2\pi$, and the phase angle of the other end is always initialized at $\theta_1 + \pi$. The degree of connectivity or topological constraint can be characterized by the filament fraction $\phi_{\mathrm{f}}$, which is defined as $\phi_{\mathrm{f}} := N_{\mathrm{f}}/N_{\mathrm{D}}$. The hard particle system with strings is further relaxed until the energy fails to change more than 1% after more than 50 iterations of relaxation. The coordinates of the center of the hard particle and strings are then used as the initial condition for the soft particle system. The soft particle system then further relaxes until none of the strings carry strain greater than the critical extension strain for rupture. After that, the actual simulation begins. Note that during the initialization, the strings experience force from the boundaries so that we can ensure all the content is inside the cell before the start of the simulation. However, during the actual simulation, the strings no longer experience force from the boundary, so we do not count the boundary repulsion multiple times when a particle is squeezing at a string at the boundary. A new Delaunay triangulation diagram is computed again at the end of the contraction, which serves as a comparison for the changes in neighboring relationships among particles. The contact number among the particles, the changes in the Delaunay diagram, potential energy, and the forces on the boundaries are recorded and used for the plot.

***A.7.7. Details on tracing neighboring relationship.*** In our analysis of the preservation of neighboring relationships, we used edges in the Delaunay diagram to trace the neighbor. However, as the particles are confined in a rectangular boundary, there are often spurious neighboring connections found in the Delaunay diagram that should not be considered true neighbors. To eliminate that, if the distance between the center of two "neighboring" particles is greater than $4 \times 1.4R_{\mathrm{c}}$, that neighboring relationship is considered spurious and eliminated from further analysis.

***A.7.8. Calculation of squeeze flow viscosity.*** As defined in Eq. 1, we calculate squeeze flow viscosity as the ratio between the normal stress in $x$-direction and the contracting strain rate in $x$-direction. For the estimation for experimental observation, based on the observed peak velocity of 0.2 m/s and half length of 0.5 mm (2), we can estimate the peak strain rate to be 400 1/sec. In Mathijssen et al. 2019, they also provided a kinematic model fitted from experimental observation, and we can also estimate the time trajectory of the contracting strain rate from that. The result is shown in Figure S6 and the peak contracting strain rate is also about 400 1/sec. The normal stress in $x$-direction can be estimated as the ratio between boundary forces and the cross-sectional area ($\sigma_{xx} \sim 10^{-6}\mathrm{N}/10^{-8}\mathrm{m}^2 \sim 100\mathrm{Pa}$). From this, we can estimate the squeeze flow viscosity of *Spirostomum ambiguum* to be roughly 0.25 Pa-sec.

In the simulation, we estimated the normal stress in $x$-direction as the ratio between the reaction force in $x$-direction and $L_y^2(t)$, and the contracting strain rate estimated as $v(t)/L_x(t)$. The squeeze flow viscosity is estimated as the ratio of the two.

***A.7.9. Validation of the models.*** As there are no previous studies on this topic that can provide a suitable validation case, we validate our code by (1) checking the force output given a known test case input, and (2) ensuring the relaxation time scale of individual particles in a small scale system matches the expected relaxation time scale listed previously.

***A.7.10. Other discussions on the biological relevance of the model.*** One concern that may arise from our model is the force that is required to be sustained at the string attachment points on the particles. In our simulation, we did not consider the detachment of strings from the particles but only consider the rupture of strings when they are stretched beyond the critical stretching strain. In the following, we show that the rupture force of the strings is comparable to the typical force magnitude of the membrane anchoring mechanism, and thus we do not have to include both mechanisms in our model.

Based on the simulation parameters described in SI Appendix Section A.7.2 and A.7.4, the force required to rupture the strings is $K_{\mathrm{A}} \times dl_{\mathrm{f}} \times \varepsilon_{\mathrm{crit}}^{\mathrm{sim}} = 2.24 \times 10^{-8}$ N. To the best of our knowledge, there is no force measurement on the rupture force of organelle contact sites in literature. We can yet still compare the rupture force in the model with the rupture force of individual integrin molecules and focal adhesions. The rupture force of an individual integrin molecule is roughly $10^{-10}$ N(21), and the typical force of a focal adhesion site is about 20 nN(22). Based on these numbers, we can thus conclude that the rupture force of strings in our model is comparable to the typical force values of membrane anchoring mechanisms and equivalent to about 200 anchoring molecules on the attachment point.

Another concern about our model is the tangential forces exerted on the vacuole surface, and whether that can displace the attachment points and un-entangle the system. Although lipid membrane has limited resistance to shear force given its fluidity, in the actual 3D case of *Spirostomum*, the attachment sites of RER on vacuoles are likely distributed across the surface, rather than just a single attachment point. Thus, tangential forces will cause stretching of the membranes, rather than just shearing at the attachment point.

***A.7.11. Sensitivity to simulation parameters.*** In all the parameters we used in the model, the measured values of interfacial tension and membrane bending rigidity tend to vary by an order of magnitude around the values chosen in SI Appendix Section A.7.4(13). In this section, we discuss the relevance of these variations to our model, and the robustness of our major conclusions despite all these variations.

As we discussed in SI Appendix Section A.7.5 and Table S1, the time scale associated with bending relaxation is $10^3$ larger than the simulation time scale. In other words, the time scale associated with the bending relaxation of the lipid bilayer membrane is much longer than the time scale associated with Spirostomum contraction (∼5 msec). That means, the bending rigidity of the lipid bilayer membrane has to be at least $10^3$ stiffer compared to the value we used to have any effect on the kinematics of the system, which is beyond the reported range. We can therefore confidently conclude that the variations in membrane bending rigidities will not change the conclusion here.

As for the interfacial tension, changing the ratio $K_3$ can indeed change the behavior of the system, but the general conclusion on the dampening effect of entangled topology does not change. In our manuscript, we originally set the ratio $K_3 = 0.25$ (Fig. S3), which is defined as the ratio between interfacial tension and area expansion modulus. In Figure S12, we show the simulation results with $K_3 = 0.025$ (low interfacial tension) and $K_3 = 2.5$ (high interfacial tension). In the low interfacial tension case (Fig. S12A-D), the particles deform more and the system contracts more given the same $\phi$ and $\phi_\mathrm{f}$. The interfacial tension and surface stretching become equally important in buffering the energy, instead of just the interfacial tension term. In the high interfacial tension case (Fig. S12E-H), the particles deform less and the system contracts less given the same $\phi$ and $\phi_\mathrm{f}$. Energy-wise, interfacial tension, particle-particle repulsion, and particle-boundary repulsion all become important in buffering energy input. Although changing the interfacial tension does affect the dominant energy terms involved, the general fact that increasing the degree of entanglement can help the system dampen the contraction kinematics and reduce the cumulative net energy input remains true.

**A.8. Table-top experiments.** To create the macroscopic counterpart of the entangled RER-vacuolar system, we used table tennis balls (KEVENZ, diameter 1.57 inch, as international standard) wrapped by sheer fabric backdrop curtains (Anteer, Crystal Organza Tulle Sheer Fabric Backdrop Curtains for Wedding Baby Shower Birthday Party Event Decor, of size 18.5 in × 23 in, with a thickness of 0.12 mm). In the topologically constrained group, the fabric was cut into interconnected stripes of roughly 2 cm thick. 45 table tennis balls were randomly attached to stripes in an interwoven fashion, and each table tennis ball was connected to 2 stripes, using super glue and protected by tapes. In the control group, the fabric was joined to form a bag, and 45 table tennis balls were put into the bag. We also applied two stripes of tape on each table tennis ball in the control group, as the presence of tape can increase the friction among balls. The bag was sealed afterward. We chose the control group to have the same amount of material, rather than control based on other parameters (such as control to be of the same convex hall) because it is closer to a biological scenario – an organism has a roughly fixed amount of resources or energy to make the same amount of RER and vacuoles, and their task is to arrange the material topologically to optimize its physical properties.

To test the ability to resist external load in two groups, the curtain in the control group was folded into the pile of table tennis balls to make a comparable height as the topologically constrained group. The samples were used for video shooting and compression testing. In the videos, two objects with different shapes and weights – a giant ruler (weight 375gm, bottom surface dimension 24 in × 2 in), and a book (weight 2450gm, bottom surface dimension 11.13 in × 8.63 in) were tested. The object was released right above the samples and the video was recorded using a smartphone.

Compression testing was done on Tensiometer Instron 5565, with compression plate of diameter 150 mm installed and a 5 kN load cell. The height of the samples was adjusted to 135 mm, and a simple compression test at a compression rate of 50mm/min with no preload was performed. The test was terminated when the displacement becomes greater than 110 mm or when the force becomes greater than 5 kN. As the compression testing inevitably destroys the sample, the compression testing was done after the video shooting, and 3 different samples were used in each group for triplicate. In addition to the compression testing on entangled and untangled samples, we also did compression testing on individual table tennis balls (simple compression test, compression rate 50 mm/min, no preload, test terminates when displacement > 35 mm or force > 5 kN) and extension testing on 2cm-thick fabric stripes (simple extension test with preload, extension rate 50 mm/min, test terminates when force drop > 40% for force > 1 kN). The reactive forces of the entangled and untangled samples were normalized by the reactive forces of the individual table tennis balls when it is compressed by 5 mm because that corresponds to a visually indisputable deformation. The peak extension force of the 2 cm fabric stripes is 37.2 ± 1.9 N (mean ± standard error), and the reactive forces of individual table tennis balls when it is compressed by 5 mm is 152.1 ± 0.7 N (mean ± standard error).

**A.9. Calculation of the spatial correlation of velocity direction.** For the simulations, the instantaneous velocity at the end of the contraction was calculated by the finite differences between the centroid coordinates of each particle at $t = 4.9999$ msec and $t = 5$ msec. For the experimental observations on contracting *Spirostomum ambiguum*, the high-speed videos (see SI Appendix Section A.2) were used. The vacuole velocity near the end of the contraction was calculated by the finite differences between the centroid coordinates of each vacuole at $t = 4$ msec and $t = 5$ msec. A larger time gap was used for experimental data to ensure the vacuoles have moved a significant distance to reduce noise. For the experimental observations on relaxed organisms, the time-lapse videos of gelatin-embedded cells (see SI Appendix Section A.2) were used. The vacuole velocity was calculated by the finite differences between the centroid coordinates of each vacuole at a time interval between 10 sec to 1 min, depending on the strength of cytoplasmic streaming in each organism. The velocities are normalized to obtain the velocity unit vector. The inner product of the velocity unit vector of vacuoles is calculated, and the self-correlations are excluded. All segmentation of vacuoles was done manually on Fiji.

### B. Perturbation experiments using ER drugs

In this section, we discuss our attempt to perturb the ER network using ER drugs. Most perturbation experiments on ER utilize ER drugs to disrupt the function and/or structure of ER. There are 2 major classes of ER drugs – the drugs that disrupt the protein folding or transport (such as tunicamycin(23) and brefeldin A(24)), and the drugs that disrupt the sarco/endoplasmic reticulum Ca2+-ATPase (SERCA) (such as thapsigargin(25) and cyclopiazonic acid(26)). Protein-folding disruption drugs commonly cause ER morphological changes such as ER dilation(27). We tested *Spirostomum ambiguum* on 1 $\mu$g/mL tunicamycin and 20 $\mu$g/mL cyclopiazonic acid. Cyclopiazonic acid almost immediately kills the cell, as the elevated calcium concentration in the cytoplasm causes tetanic contraction followed by cell membrane dissociation. On the other hand, tunicamycin did not disrupt the ER-vacuole entanglement morphology under confocal microscopy (see Movie S13). This suggests, despite the potential changes in ER morphology under electron microscopy, the changes are not large enough to disrupt the ER-vacuole entanglement structure to perturb the stalling mechanism.

It should also be noted that ER plays a dual role in the contraction and the stalling process. It is the storage of calcium, and the rapid release and efficient delivery of calcium from ER is necessary for the ultrafast contraction to happen in the first place. It is currently unclear whether the ER-vacuole topology that we observed has any implications for efficient signaling transduction inside the cell, and disrupting ER networks can also disrupt the calcium release. Therefore, even if the drug causes any changes in contraction kinematics, it will be hard to clearly interpret the results. The disruption of ER-vacuole entangled structures would likely require a massive disruption of the ER, and that could have killed the cell before any changes in the stalling mechanism could be observed.

### C. Further discussions on volume fractions of vacuoles

We estimated the volume fraction of vacuoles using confocal microscopy data, and the estimated volume fraction is 65.0% (n=3, std = 6.3%). This is very close to the critical volume fraction (63.9%) for jamming transition in 3D hard monodisperse spheres(28). However, note that polydisperse particles (as shown in Figure S9) actually require a larger critical volume fraction to have jamming transition. For polydisperse particles with power law distribution, the critical volume fraction can go above 90% (29). It is thus not as useful to deduce whether the cytoplasm is jammed by looking at the vacuole volume fraction, and the velocity direction spatial correlation described in the main text is a better way to identify jamming transition.

### D. Discussions on lipid droplets

In this manuscript, we focus on the entangled architecture between RER and vacuoles, and there may be concerns about other organelles also forming the entangled architecture. In our TEM images, there are not many lipid droplets that can form mechanical contact with the vacuoles. We have also done confocal microscopy on *Spirostomum ambiguum* stained with Nile red, but only a few lipid droplets are identified and are outnumbered by mitochondria (Fig. S10). The only dominant organelles that are present in large quantities in *Spirostomum ambiguum* are the vacuolar meshwork, RER, and mitochondria, the ones that we show in Figure 2B in the main text.

### E. The discrepancy of vacuole deformations between simulations and experiments

In Figure S4, we show that our model allows us to follow the deformation of each particle just like in the experiments. However, we acknowledge that the magnitude of vacuole deformations is different from experiments. This likely comes from the following reasons. First, the 3D ER fenestrated membrane sheets are simplified into 2D strings, which have much less capability to “cage” the vacuoles and force their deformation. Second, there might be distributed attractive forces between RER and vacuoles, either through the Casimir effect (described in Section "The nature of RER-vacuole contact sites" in the main text) or protein linkage. The attractive force between the two can make vacuoles conforming to the deformations of RER more energetically favorable. However, in our model, we did not account for the distributed attractive forces throughout the RER and vacuolar surfaces. Third, the interfacial tension of vacuoles has more variations in reported values compared to other parameters. In Figure S12, we indeed observe that lowering the interfacial tension of particles in simulation makes the deformation of particles greater.

That being said, we present the cleanest reduced order system in order to demonstrate that just the entanglement effect alone is sufficient to transform the mechanical properties of the cytoplasm, and the vacuoles do not have to be drastically deformed to have strain-induced jamming transition. We thus believe this discrepancy between predicted and observed aspect ratio changes in vacuoles does not change the main conclusion of this work.

### F. Negligible role of heating

The discussion on energy dissipation also raises the concern about heating. However, some simple calculations will show that the energy associated with the contraction process will not cause significant heating inside the cell. We can estimate the total work during the contraction as $W = F\Delta x \sim FL \sim 10^{-9}$J. Assuming that the density and heat capacity of the cell is similar to water, we can estimate the temperature changes in the cytoplasm after one cycle of contraction to be about $\Delta T \sim 2 \times 10^{-5}$ °C. It is thus safe to conclude that heating is not likely a major issue for *Spirostomum*.

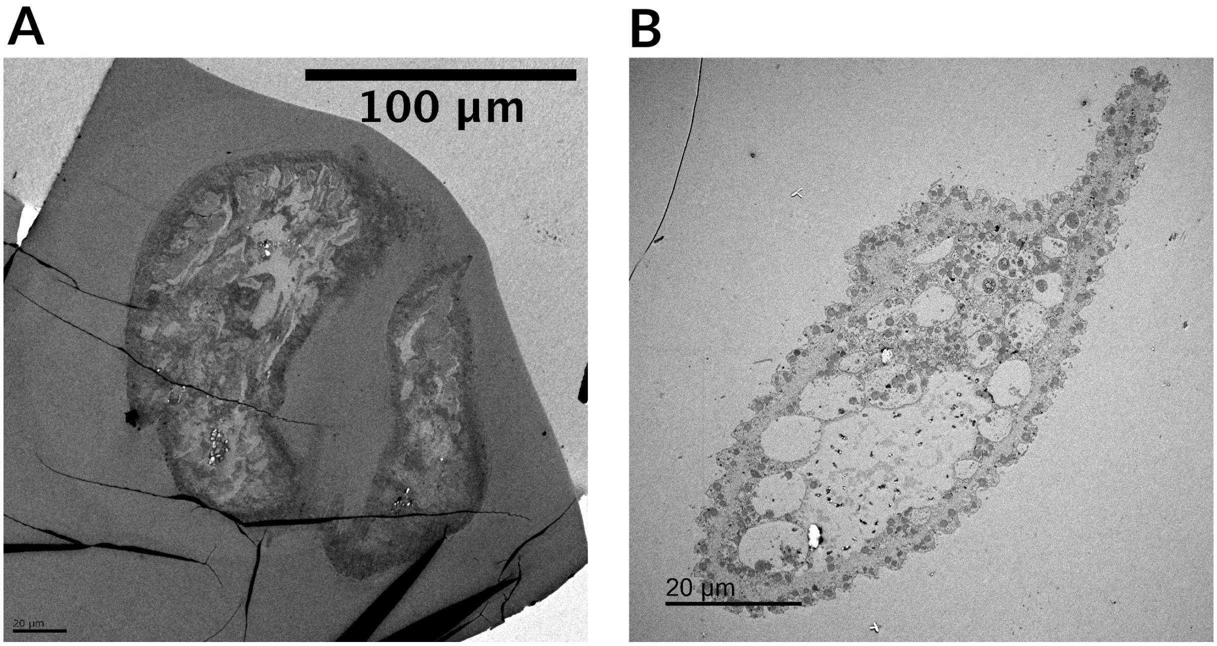

**Fig. S1.** TEM image of the entire cross-section of (A)contracted and (B)relaxed *Spirostomum ambiguum*. The cytoplasm is vacuolated in both cases.

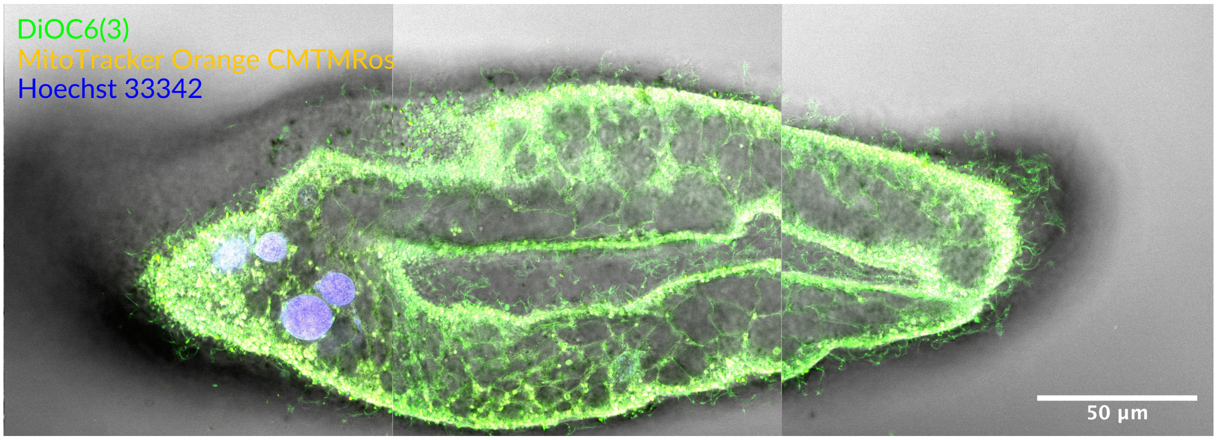

**Fig. S2.** Confocal microscopy of *Spirostomum ambiguum*, with staining of DiOC6(3), Mitotracker Orange CMTMRos, and Hoechst 33342. We can see the wrapping relationship between ER and vacuoles span through the entire organism.

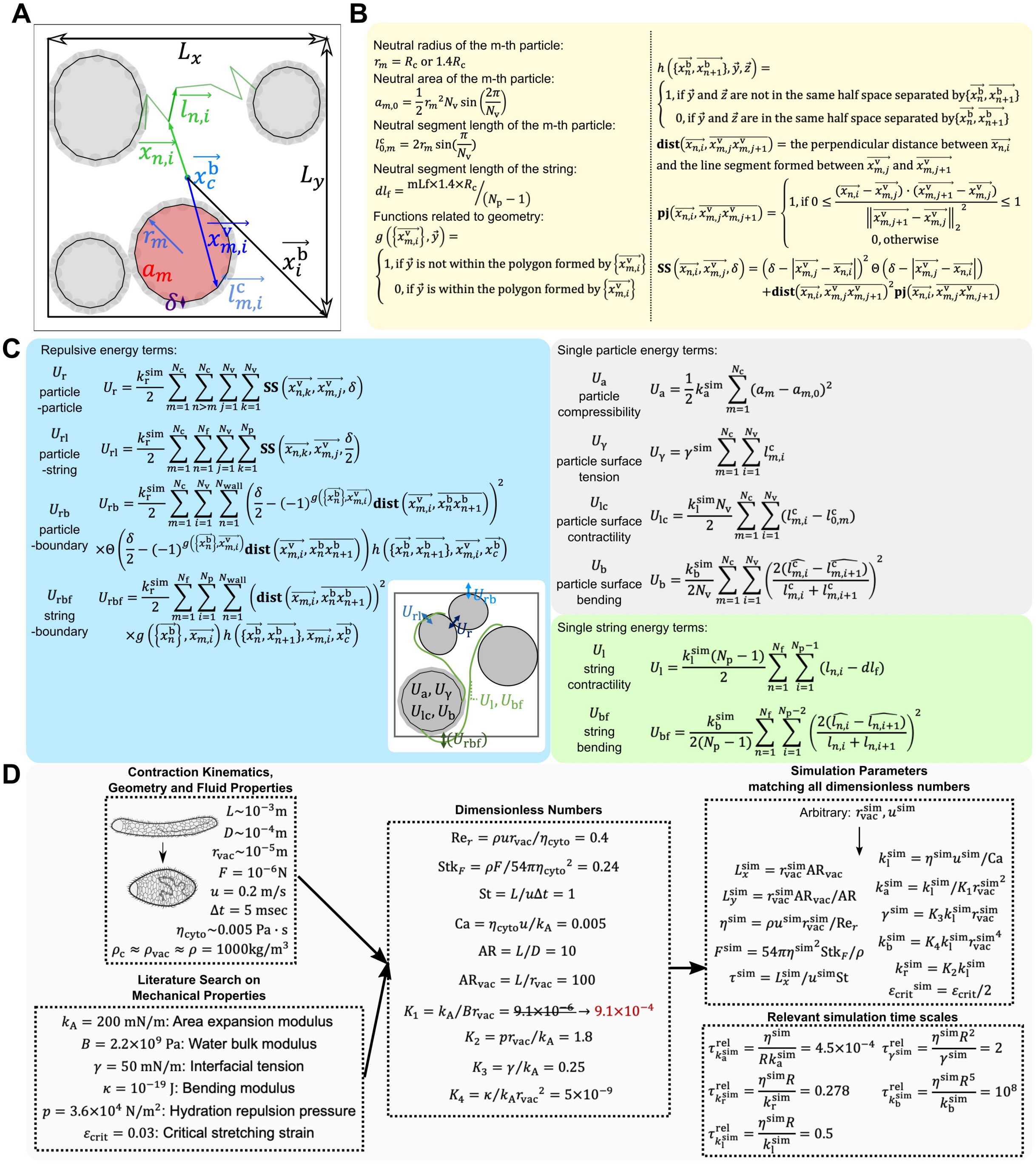

**Fig. S3.** Detailed description of the model. (A) A schematic diagram illustrating the relevant definition in the model. See SI Appendix Section A.7.1-A.7.2 for the descriptions. Note that the particles are deformable and bidisperse. (B) Relevant geometrical definitions to the simulation. (C) Energy terms defined in the model, with the inset diagram showing each term. Note that $\hat{(\cdot)} = \vec{(\cdot)}/|\vec{(\cdot)}|$. (D) Detailed process to determine the parameters. Known parameters from contraction kinematics, geometry and fluid properties (including the length of the organism $L$, the diameter of the organism $D$, the characteristic radius of the vacuole $r_{\mathrm{vac}}$, the boundary forces $F$(11)), characteristic velocity $u$(2), characteristic time scale $\delta t$(2), cytoplasm viscosity(12) (measurement in other protists), fluid density $\rho$) are combined with known mechanical properties of lipid bilayer membrane (area expansion modulus $k_{\mathrm{A}}$(13), water bulk modulus $B$(14), interfacial tension $\gamma$ (13), bending modulus $\kappa$(13), hydration repulsion pressure of lipid bilayer membrane $p_0$(15), and the critical stretching strain of lipid bilayer membrane before rupture $\varepsilon_{\mathrm{crit}}$ (16)) to generate 10 dimensionless numbers. These dimensionless numbers include: Reynolds number based on vacuole radius $\mathrm{Re}_r$, Stokes number based on external force $\mathrm{Stk}_F$, Strouhal number St, capillary number based on membrane stretching Ca, aspect ratio of the cell AR, aspect ratio between the length of the cell and the radius of the vacuole $\mathrm{AR}_{\mathrm{vac}}$, and the four relative strength ratio between the 5 membrane mechanical properties $K_1$, $K_2$, $K_3$, and $K_4$, following the definition in Boromand et al(7). The dimensionless number $K_1$ is multiplied by 100 as the systems of ODE are too stiff. This operation is equivalent to assuming that the bulk modulus of water is 100 times smaller, but it is still sufficiently large to ensure area conservation of the 2D vacuoles. The critical strain for rupturing string is divided by 2, as we have to convert the area expansion strain of lipid bilayer membranes in three-dimension into the extension strain of strings in two-dimension. In the simulation, the radius of the vacuole and characteristic velocity can be arbitrarily set (which determines the absolute value of length scale and time scale), and the remaining parameters are set by matching the 10 dimensionless numbers. Five relevant relaxation time scales in the simulation are also listed, and we determined the time step according to these time scales (see SI Appendix Section A.7.5).

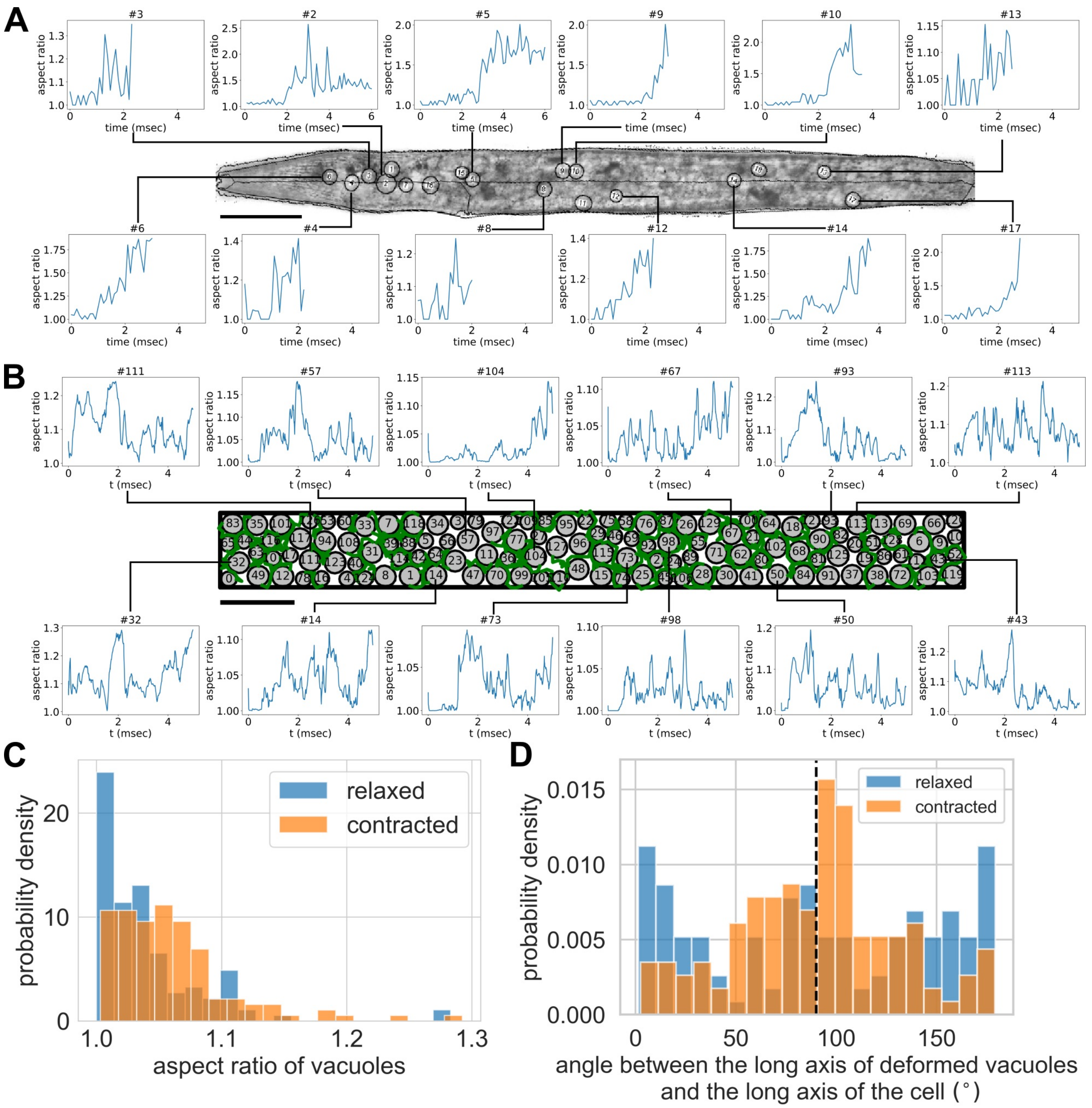

**Fig. S4.** Comparison of vacuole deformation between (A) experiment and (B) simulations. (A) High speed video of the contraction process along with the shape changes of vacuoles are shown. Most vacuoles cannot be tracked for the entire duration of contraction because the twisting motion of the organism brings the vacuoles away from the focal plane. The onset of shape deformation of each vacuole depends on its axial position. Scale bar = 100 $\mu$m. (B) Simulation with volume fraction $\phi = 0.79$ and filament fraction $\phi_{\mathrm{f}} = 0.268$, with the time series changes of the aspect ratio of selected vacuoles. Scale bar = 100$\mu$m. (C) The histogram of the aspect ratio of deforming particles in relaxed and contracted organisms in simulations. (relaxed: mean = 1.040, std = 0.044, 130 particles; contracted: mean = 1.059, std = 0.047, 130 particles.)(D) The histogram of the angle between the long axis of the particles and the long axis of the cell body before and after the contraction depicts the deformation is normal to the external load. (relaxed: mean = 91.3$^\circ$), std = 58.0$^\circ$, 130 particles; contracted: mean = 89.9$^\circ$, std = 40.1$^\circ$, 130 particles. (See SI Appendix, Section E for further discussion.)

 

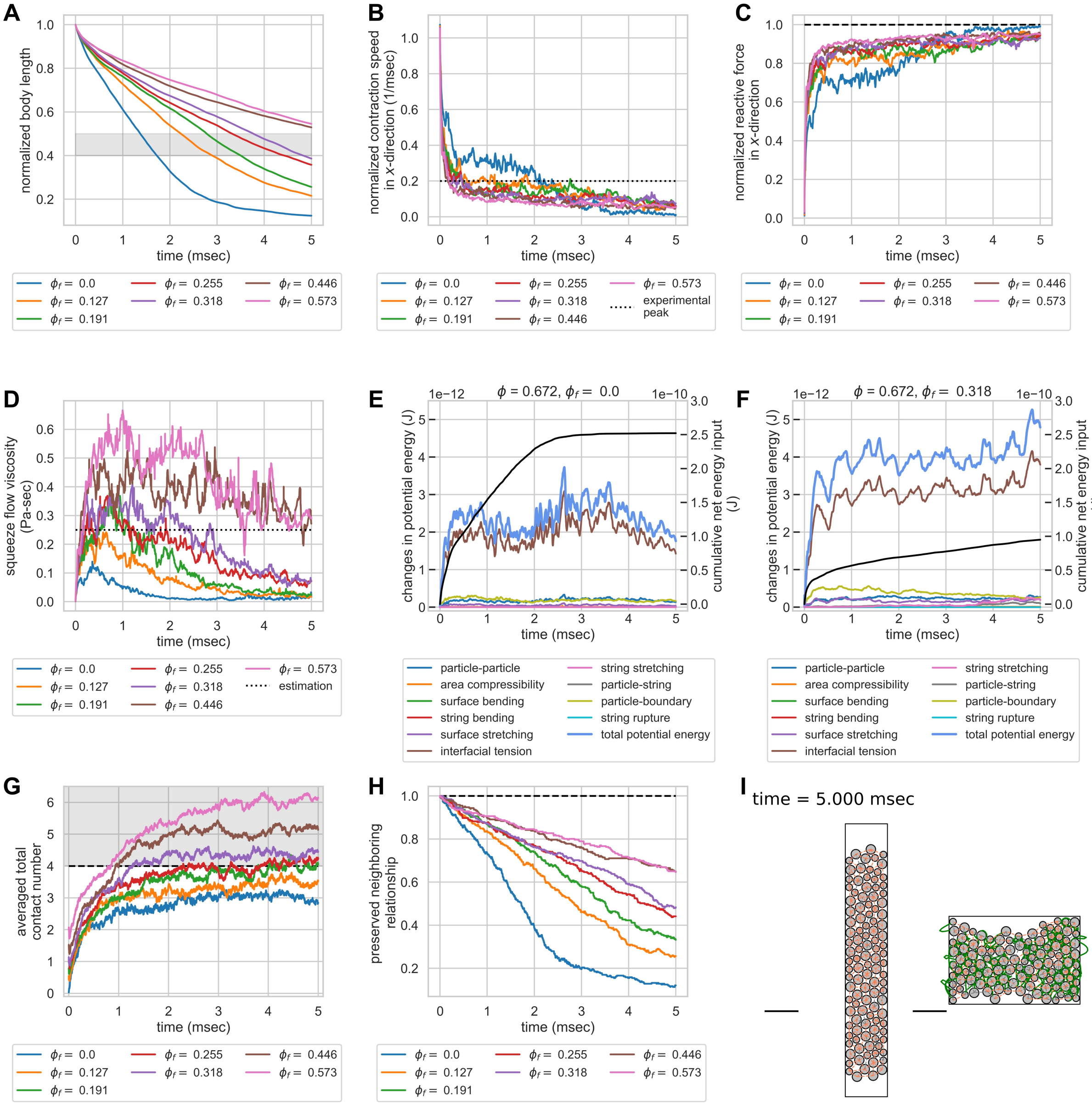

**Fig. S5.** Geometric constraints turn an entangled particle system into a topological damper in constant force simulation. Systems with the same $\phi = 0.67$ but different $\phi_\mathrm{f}$ were compressed by a constant boundary force. (A) Normalized body length as a function of time. The gray zone indicates the experimentally observed final body length after contraction (40%-50%)(30). The system with $\phi_\mathrm{f} = 0.318$ roughly falls within the range. (B) Normalized contraction rate in the $x$ direction as a function of time. The dashed line indicates the experimentally observed peak velocity of 0.2 m/sec (2). Systems with $\phi_\mathrm{f} \geq 0.255$ can rapidly dampen the velocity to be below the experimentally observed peak velocity, while systems with lower $\phi_\mathrm{f}$ fail to dampen the velocity, and maintain their contraction velocity at peak level for roughly 2 msec. (C) Time series changes of the reactive forces of the system in $x$-direction normalized with respect to the external load. Increasing topological constraints increases the ability of the system to counteract external forces, which explains the rapid slowdown in contraction kinematics. The dashed horizontal line indicates the magnitude of external load, normalized to 1. (D) Squeeze flow viscosity of the system as a function of time. The dotted line is the estimation of actual organisms based on the peak strain rate and normal stress in $x$-direction (0.25 Pa-sec). Systems with $\phi_\mathrm{f} \geq 0.255$ maintain their squeeze flow viscosity at a level comparable to estimation from experiments, while systems with $\phi_\mathrm{f} < 0.255$ fail to maintain their squeeze flow viscosity despite the initial peak. (E)&(F) Energy budget of 2 systems with the same $\phi = 0.67$ but $\phi_\mathrm{f}$ of 0.0 and 0.318. The cumulative net energy input is plotted in black line, while other colored lines are different energy terms in the system. The system with no entanglement has a larger cumulative net energy input, yet the system with entanglement is able to store more potential energy. Note that the majority of the energy contribution comes from the interfacial tension terms of the particles, and the contributions from strings are minimal. The sacrificial bond effect from string rupture is also negligible. (G) Averaged total contact number of the particles as a function of time for systems with different $\phi_\mathrm{f}$. Systems with $\phi_\mathrm{f} \geq 0.318$ can undergo strain-induced jamming transition as the averaged total contact number is greater than 4. (H) The preserved neighbor of the system as a function of time. Increasing topological constraints helps preserve the neighboring relationship after a contraction event. (I) Preserved neighbor after a single contraction event for two systems. The neighbors are identified as the edges on the Delaunay diagram, and the list of neighboring particles is compared before and after contraction. New neighbors are colored in pink and preserved neighbors are colored in gray. The figure on the left-hand side has $\phi_\mathrm{f} = 0$, and 12.1% of the neighbors are preserved. The figure on the right-hand side has $\phi_\mathrm{f} = 0.318$, and 48.3% of the neighbors are preserved. The volume fraction of the system is 67%. Scale bar = 100 $\mu$m.

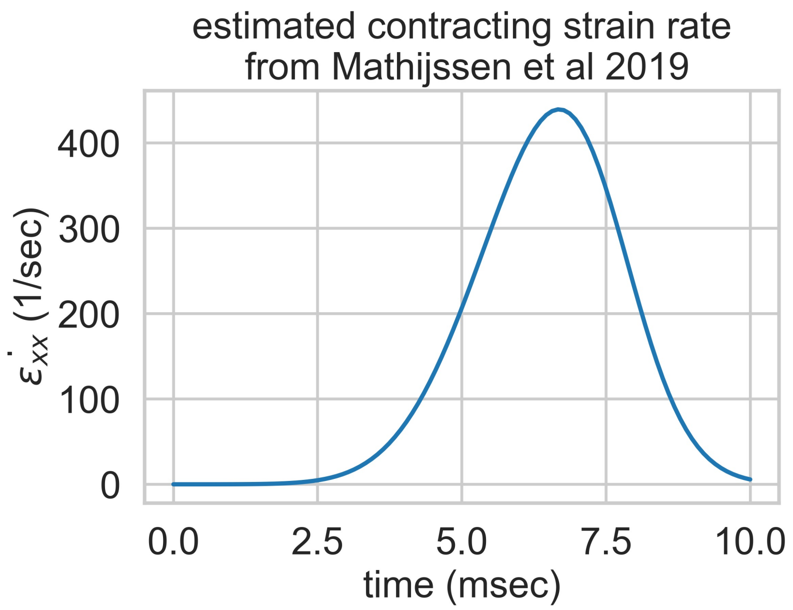

**Fig. S6.** Contracting strain rate estimation using the empirical kinematics model reported in Mathijssen et al. 2019(2). The empirical formula of organism length as a function of time can be described as $L(t) = L_0 \left(1 - \frac{f_c}{2}\left(\mathrm{erf}\,\frac{\tilde{\tau_p}}{\sqrt{2}\tilde{\tau_c}} - \mathrm{erf}\,\frac{\tilde{\tau_p} - t}{\sqrt{2}\tilde{\tau_c}}\right)\right)$, where initial body length $L_0$ = 1 mm, fraction of length contraction $f_c$ = 0.5, perception time $\tau_p$ = 3.5 msec, contraction duration $\tau_c$ = 5 msec, with $\tilde{\tau_p} = \tau_p + \tau_c/2$ and $\tilde{\tau_c} = \tau_c/4$. Contraction velocity $v(t) = \frac{\mathrm{d}L}{\mathrm{d}t}$. The contracting strain rate is estimated as $\dot{\varepsilon_{xx}}(t) = v(t)/L(t)$.

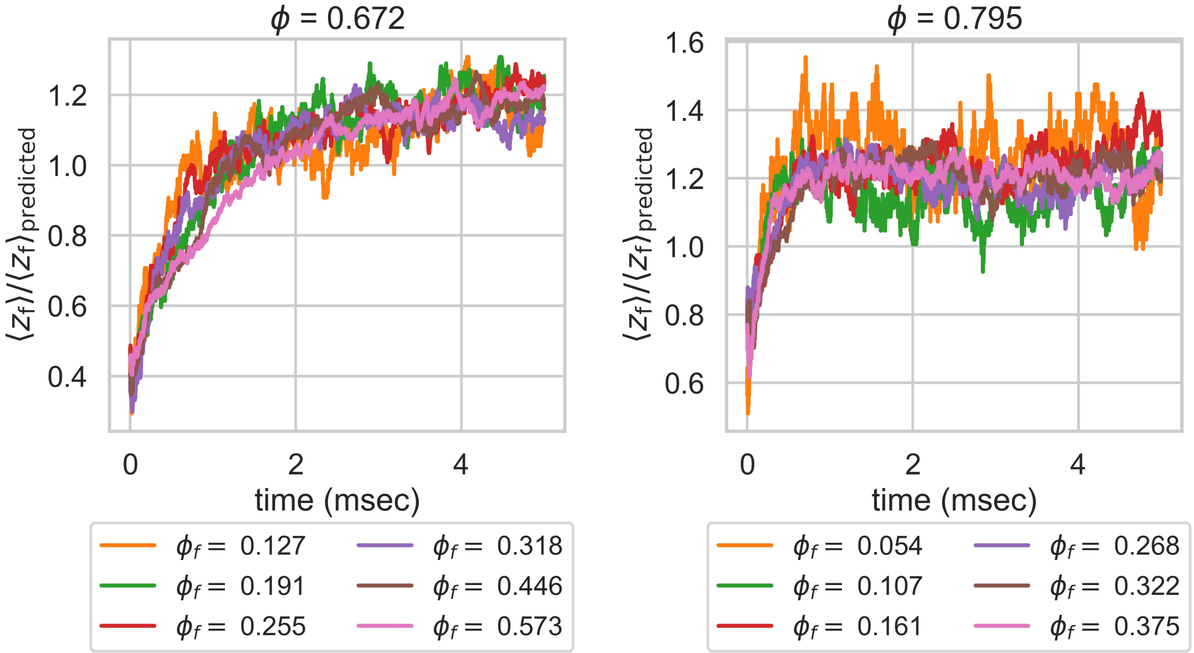

**Fig. S7.** Comparison between the predicted mean contact number from strings ($\langle z_{\mathrm{f}} \rangle_{\mathrm{predicted}}$) from probability theory and the observed mean contact number from strings in simulation ($\langle z_{\mathrm{f}} \rangle$). The figure shows the time series changes of the ratio between the two contact numbers for systems with $\phi$ of 0.672 (left) and 0.795 (right). The time series curves for systems with different $\phi_{\mathrm{f}}$ collapse to a single curve and reach the same plateau value of 1.2. Systems with larger volume fractions reach the plateau value faster. The constant ratio of 1.2 probably depends on the nature of the strain itself (e.g. squeeze strain v.s. shear strain).

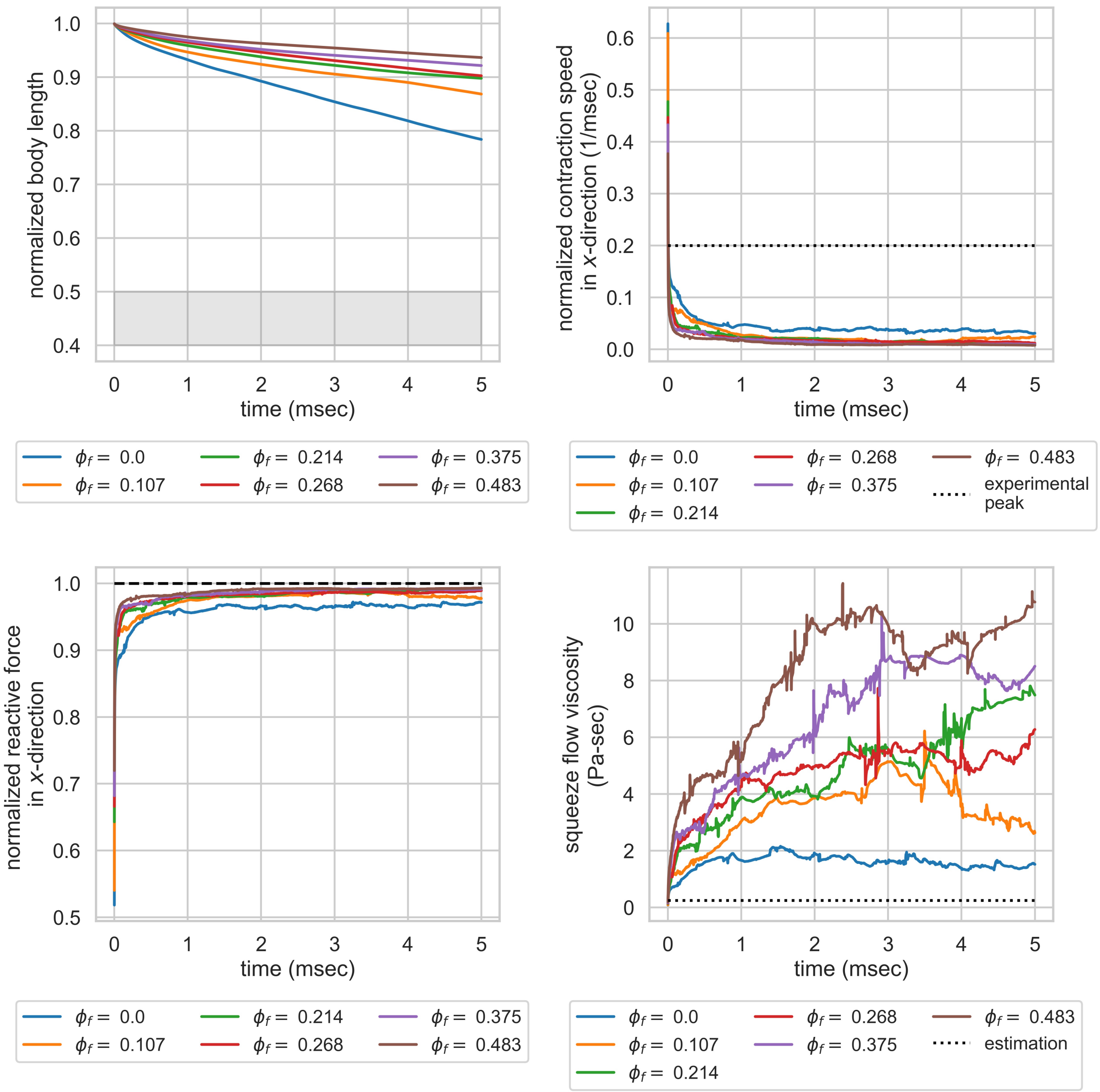

**Fig. S8.** Simulations with high viscosity (cytoplasm viscosity = 0.05 Pa-sec). The systems have $\phi = 0.79$ with varying $\phi_{\mathrm{f}}$. Even for the systems with $\phi_{\mathrm{f}} = 0$, the final normalized body length is still way above the experimentally observed final body length. The contraction speed plummets to levels way below experimental observation, and the normalized reactive forces are high. The squeeze flow viscosity of the systems is also way above the estimation from experiments. These results suggest that the cytoplasm viscosity of *Spirostomum ambiguum* should be on the lower end of the reported range from other protists.

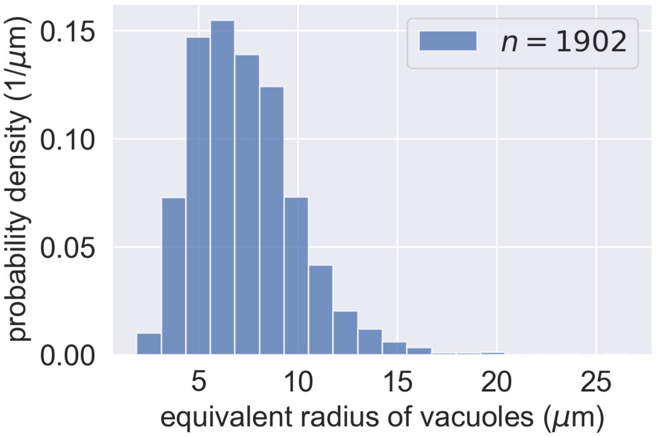

**Fig. S9.** Size distribution of vacuoles in *Spirostomum ambiguum*. Statistics are from 1902 vacuoles from 3 relaxed organisms. (mean = 7.33 $\mu$m, std = 2.66 $\mu$m.)

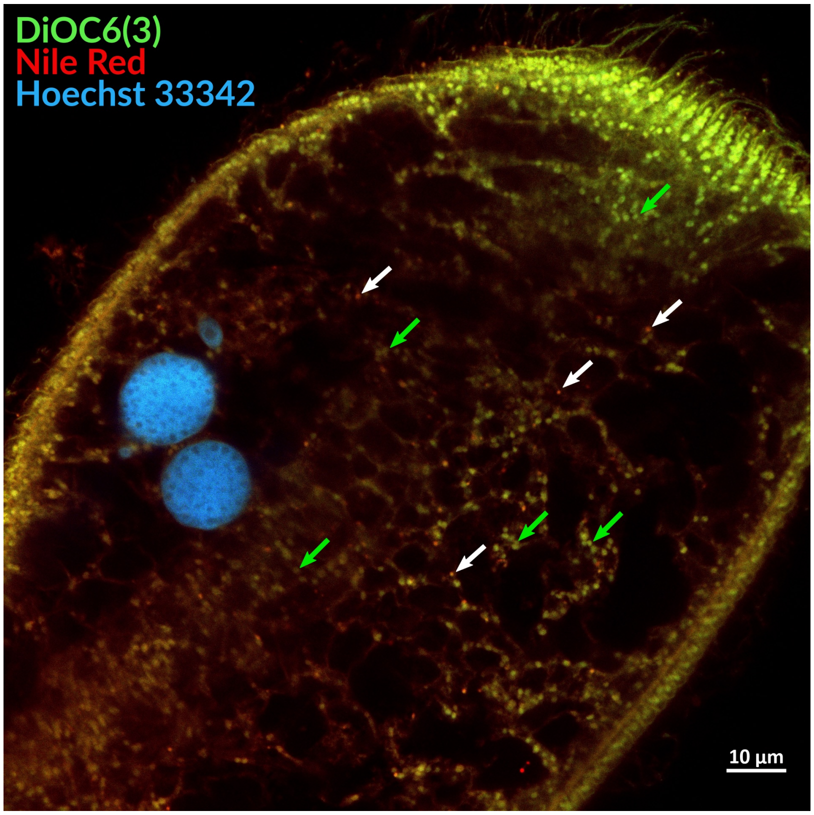

**Fig. S10.** Confocal microscopy of *Spirostomum ambiguum*, with staining of Nile red, DiOC6(3), and Hoechst 33342. Cell membrane, ER (identified by DiOC6(3)), mitochondria (identified by DiOC6(3)), and nuclear membrane (identified by Hoechst 33342) are stained. Only a few lipid droplets are identified (white arrows) and are outnumbered by mitochondria (green arrows). The vacuoles are not lipid droplets.

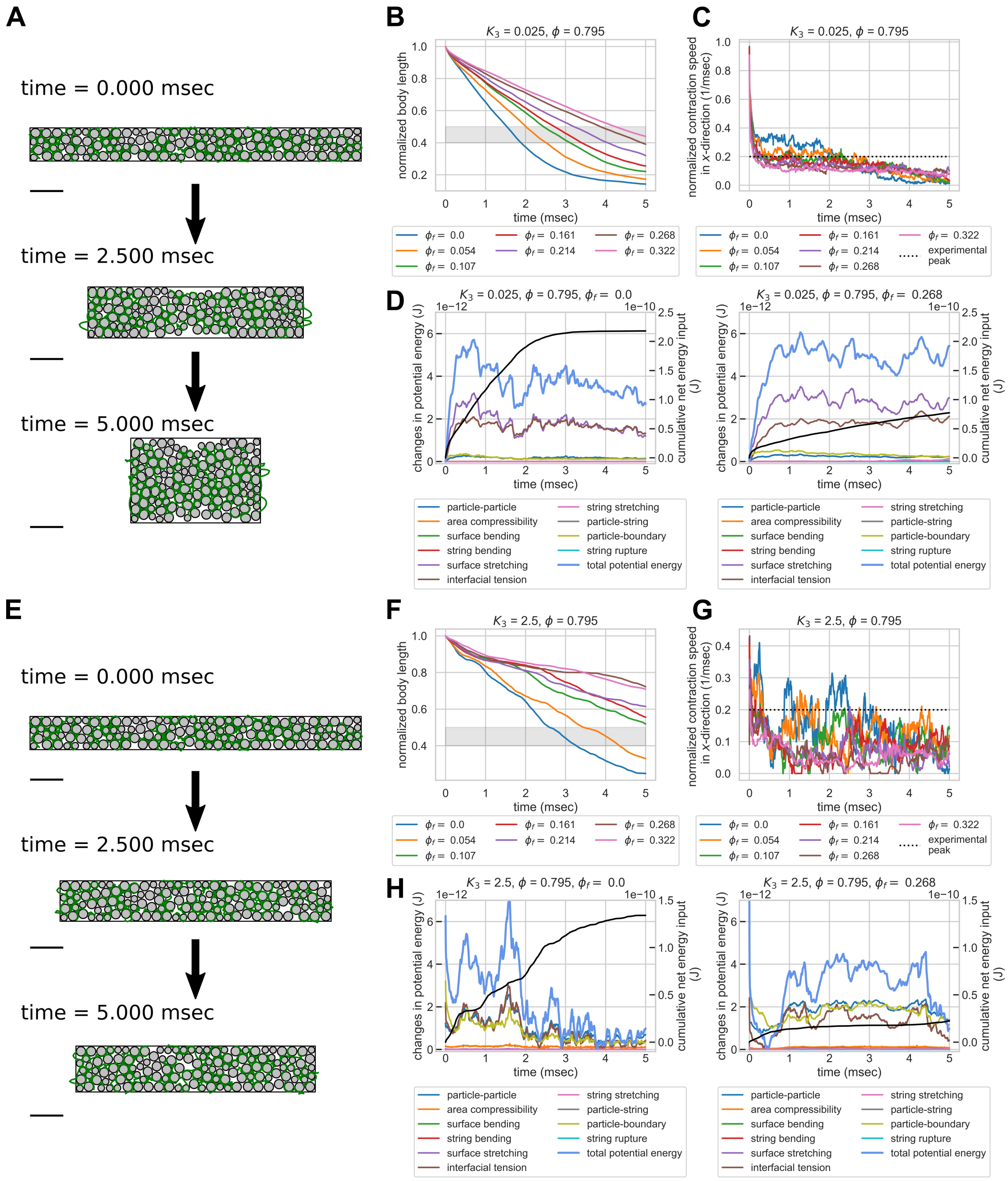

**Fig. S11.** Topological damping is not sensitive to changes in interfacial tension term of the model. We change the model parameter $K_3$ (described in Figure S3) from 0.25 (baseline) to 0.025 (low interfacial tension, (A)-(D)) and 2.5 (high interfacial tension, (E)-(H)) and see its effect on model predictions. (A)&(E) shows three snapshots of the simulated contraction process of a system with $\phi = 0.795$ and $\phi_\mathrm{f} = 0.268$. The system with low interfacial tension (A) shows faster contraction and greater vacuolar deformation, while the system with high interfacial tension (E) shows slower contraction and less vacuolar deformation. Scale bar = 100 $\mu$m. (B) Normalized body length as a function of time ($K_3 = 0.025$). The gray zone indicates the experimentally observed final body length after contraction (40%-50%)(30). The system with $\phi_\mathrm{f} = 0.322$ roughly falls within the range. (C) Normalized contraction rate in the $x$ direction as a function of time ($K_3 = 0.025$). The dashed line indicates the experimentally observed peak velocity of 0.2 m/sec (2). Systems with $\phi_\mathrm{f} \geq 0.268$ can rapidly dampen the velocity to be below the experimentally observed peak velocity, while systems with lower $\phi_\mathrm{f}$ fail to dampen the velocity, and maintain their contraction velocity at peak level for roughly 2 msec. (D) Energy budget of 2 systems with the same $\phi = 0.795$ and $K_3 = 0.025$ but $\phi_\mathrm{f}$ of 0.0 and 0.268. The cumulative net energy input is plotted in black line, while other colored lines are different energy terms in the system. The system with no entanglement has a larger cumulative net energy input, yet the system with entanglement is able to store more potential energy. Different from the case in $K_3 = 0.25$, the majority of the energy contribution comes from the interfacial tension and surface stretching terms of the particles. The contributions from strings and the sacrificial bond effect from string rupture are both minimal. (F) Normalized body length as a function of time ($K_3 = 2.5$). The gray zone indicates the experimentally observed final body length after contraction (40%-50%)(30). The system with $\phi_\mathrm{f} = 0.107$ roughly falls within the range. (G) Normalized contraction rate in the $x$ direction as a function of time ($K_3 = 2.5$). The dashed line indicates the experimentally observed peak velocity of 0.2 m/sec (2). Systems with $\phi_\mathrm{f} \geq 0.161$ can rapidly dampen the velocity to be below the experimentally observed peak velocity, while systems with lower $\phi_\mathrm{f}$ fail to dampen the velocity, and maintain their contraction velocity at peak level for roughly 2-3 msec. (H) Energy budget of 2 systems with the same $\phi = 0.795$ and $K_3 = 2.5$ but $\phi_\mathrm{f}$ of 0.0 and 0.268. The cumulative net energy input is plotted in black line, while other colored lines are different energy terms in the system. The system with no entanglement has a larger cumulative net energy input, yet the system with entanglement is able to store more potential energy. Different from the case in $K_3 = 0.25$, the majority of the energy contribution comes from the interfacial tension, particle-particle repulsion, and particle-boundary repulsion. The contributions from strings and the sacrificial bond effect from string rupture are both minimal.

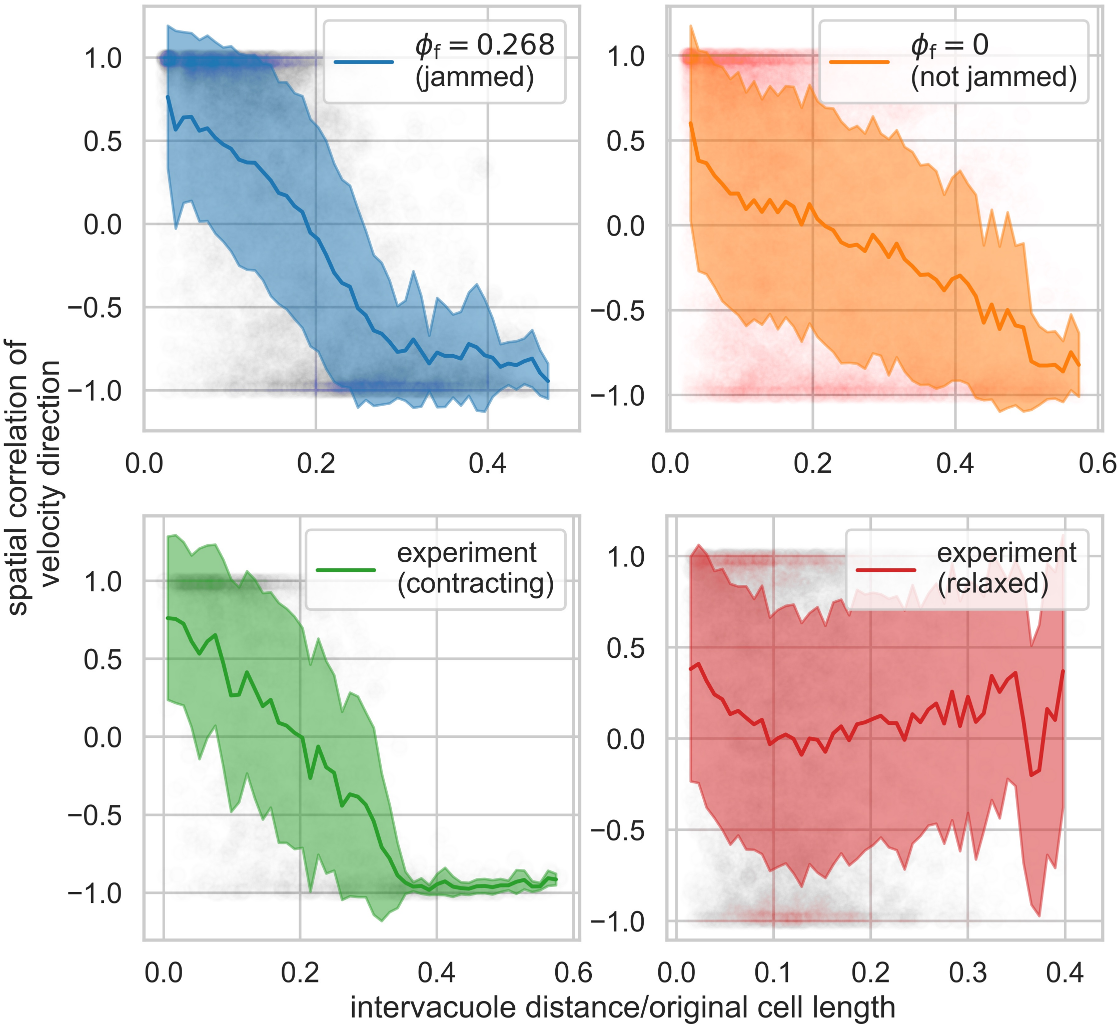

**Fig. S12.** Spatial correlation of velocity direction as a function of normalized intervacuole distance (mean $\pm$ standard deviation), overlaid with raw data. Jammed cases in simulations and contracting organisms (3 organisms, total 118 vacuoles) exhibit slower decay in spatial correlation, with negative correlation when normalized distance exceeds 0.3 due to confinement. The unjammed case shows a faster decay in spatial correlation and a negative correlation at larger distances. Relaxed organisms (3 organisms, total 182 vacuoles) show fast decay in spatial correlation at short distances and nearly-zero correlation as the normalized distance goes beyond 0.1. These indicate that the vacuolar meshwork is not jammed in relaxed organisms but jammed in contracting organisms, constituting a strain-induced jamming transition. The data is the same as is shown in Figure 5G in the main text.

**Movie S1. High speed video of the contraction process of *Spirostomum ambiguum.* The video was acquired at 10kfps and is slowed down 250×. We can clearly see the deformation of vacuoles during the contraction process.**

**Movie S2. High speed video of the contraction process of another *Spirostomum ambiguum.* The video was acquired at 10kfps and is slowed down 500×. We can clearly see the deformation of vacuoles during the contraction process.**

**Movie S3. Confocal microscopy of *Spirostomum ambiguum*, with staining of DiOC6(3), Mitotracker Orange CMTMRos, and Hoechst 33342. The interval in z was 2.5 $\mu$m, and the total span in the z-direction was 67.5 $\mu$m. We can see the wrapping relationship between ER and vacuoles span through the entire organism.**

**Movie S4. Confocal microscopy of *Spirostomum ambiguum* (same in Figure 2G), with staining of DiOC6(3), Mitotracker Orange CMTMRos, and Hoechst 33342. The interval in z was 0.36 $\mu$m, and the total span in the z-direction was 26.49 $\mu$m.**

**Movie S5. Zoomed-in video of the confocal image of the endoplasmic reticulum of *Spirostomum ambiguum* (same in Figure 2G), stained with DiOC6(3). The interval in z was 0.36 $\mu$m, and the total span in the z-direction was 26.49 $\mu$m. The ER forms a fenestrated sheet surrounding the vacuoles, confirming the picture we saw in TEM.**

**Movie S6. Simulated contraction process of an entangled soft particle system with volume fraction $\phi = 0.79$ and filament fraction $\phi_f = 0$. Particles marked as red are jammed. We can see that in an untangled system, the contraction kinematics is fast and there is a massive rearrangement of particles during the contraction process. Scale bar 100 $\mu$m.**

**Movie S7. Simulated contraction process of an entangled soft particle system with $\phi = 0.79$ and $\phi_f = 0.268$. We can see that in the entangled system, the contraction kinematics is greatly slowed down compared to video S6, and the particles were jammed with less rearrangement. Scale bar 100 $\mu$m.**

**Movie S8. Table-top experiments using table tennis balls and curtain fabrics demonstrate the differences in mechanical properties between entangled (right) and untangled (left) systems. The two systems have the same number of table tennis balls and the same amount of fabrics. In this experiment, a 375gm giant ruler was placed on top of the two samples. The entangled system has a better ability to resist the load.**

**Movie S9. Table-top experiments using table tennis balls and curtain fabrics demonstrate the differences in mechanical properties between entangled (right) and untangled (left) systems. In this experiment, a 2450gm book was placed on top of the two samples. The entangled system has a better ability to resist the load.**

**Movie S10. Compression testing on the untangled system composed of 45 table tennis balls and backdrop curtains. As the volume fraction of the system is below the critical value of jamming transition, the reactive forces decrease when the system relaxes.**

**Movie S11. Compression testing on the entangled system composed of 45 table tennis balls and backdrop curtains. In the middle of the testing, the sound of the fabrics tearing is audible, and that corresponds to a drop in reactive forces before the compression reaches 95mm. The final pop sound comes from the break of a table tennis ball.**

**Movie S12. Time lapse video of the relaxed *Spirostomum ambiguum* embedded in 2% gelatin. The video was acquired at 0.1 fps, playing 300× faster. We can clearly see the cytoplasmic streaming of vacuoles, indicating that the vacuolar meshwork before contraction is not jammed.**

**Movie S13. Confocal microscopy of *Spirostomum ambiguum* treated with 1 $\mu$g/mL tunicamycin (an ER drug) overnight, with staining of DiOC6(3), MitoSpy Orange CMTMRos, and Hoechst 33342. The interval in z was 0.32 $\mu$m, and the total span in the z-direction was 22.72 $\mu$m. We can see the wrapping relationship between ER and vacuoles do not change.**

### References

1. PJ Hummer, Culturing & Using Protozoans in the Laboratory. *Am. Biol. Teach.* **55**, 357–360 (1993).
2. AJ Mathijssen, J Culver, MS Bhamla, M Prakash, Collective intercellular communication through ultra-fast hydrodynamic trigger waves. *Nature* **571**, 560–564 (2019).
3. N Boggs, Jr, Comparative studies onspirostomum: Silver impregnation of three species. *The J. Protozool.* **12**, 603–606 (1965).
4. M Schliwa, J Van Blerkom, Structural interaction of cytoskeletal components. *J. Cell Biol.* **90**, 222–235 (1981).
5. L Carter, Ionic Regulation in the Ciliate Spirostomum Ambiguum. *J. Exp. Biol.* **34**, 71–84 (1957).
6. P Singhal, et al., Evaluation of Histomorphometric Changes in Tissue Architecture in Relation to Alteration in Fixation Protocol – An Invitro Study. *J. Clin. Diagn. Res. : JCDR* **10**, ZC28 (2016).
7. A Boromand, A Signoriello, F Ye, CS O'Hern, MD Shattuck, Jamming of Deformable Polygons. *Phys. Rev. Lett.* **121**, 248003 (2018).
8. K Vanderwerf, W Jin, MD Shattuck, CS O'Hern, Hypostatic jammed packings of frictionless nonspherical particles. *Phys. review. E* **97**, 012909 (2018).
9. DJ Koeze, D Vågberg, BB Tjoa, BP Tighe, Mapping the jamming transition of bidisperse mixtures. *EPL* **113**, 54001 (2016).
10. GJ Gao, J Bławzdziewicz, CS O'Hern, Frequency distribution of mechanically stable disk packings. *Phys. Rev. E - Stat. Nonlinear, Soft Matter Phys.* **74**, 061304 (2006).
11. RB Hawkes, DV Holberton, Myonemal contraction of spirostomum. II. Some mechanical properties of the contractile apparatus. *J. Cell. Physiol.* **85**, 595–602 (1975).
12. SS Rogers, TA Waigh, JR Lu, Intracellular Microrheology of Motile Amoeba proteus. *Biophys. J.* **94**, 3322 (2008).
13. R Dimova, CM Marques, eds., *The giant vesicle book.* (CRC Press, Taylor & Francis Group), pp. 85, 284–302 (2019).
14. RA Fine, FJ Millero, Compressibility of water as a function of temperature and pressure. *The J. Chem. Phys.* **59**, 5529 (1973).
15. SA Simon, CA Fink, AK Kenworthy, TJ McIntosh, The hydration pressure between lipid bilayers. Comparison of measurements using x-ray diffraction and calorimetry. *Biophys. J.* **59**, 538–546 (1991).
16. LA Bagatolli, D Needham, Quantitative optical microscopy and micromanipulation studies on the lipid bilayer membranes of giant unilamellar vesicles. *Chem. Phys. Lipids* **181**, 99–120 (2014).
17. RHJ Brown, The Protoplasmic Viscosity of Paramecium. *J. Exp. Biol.* **17**, 317–324 (1940).
18. CF Curtiss, JO Hirschfelder, Integration of Stiff Equations. *Proc. Natl. Acad. Sci.* **38**, 235–243 (1952).
19. HP Langtangen, L Wang, Odespy software package (2015) https://github.com/hplgit/odespy.
20. B Delaunay, Sur la sphère vide. a la mémoire de georges voronoï. *Bull. de l'Académie des Sci. de l'URSS. Cl. des sciences mathématiques et na* pp. 793–800 (1934).
21. F Li, SD Redick, HP Erickson, VT Moy, Force Measurements of the $\alpha5\beta1$ Integrin–Fibronectin Interaction. *Biophys. J.* **84**, 1252–1262 (2003).
22. US Schwarz, et al., Calculation of Forces at Focal Adhesions from Elastic Substrate Data: The Effect of Localized Force and the Need for Regularization. *Biophys. J.* **83**, 1380–1394 (2002).
23. P Guha, E Kaptan, P Gade, DV Kalvakolanu, H Ahmed, Tunicamycin induced endoplasmic reticulum stress promotes apoptosis of prostate cancer cells by activating mTORC1. *Oncotarget* **8**, 68191–68207 (2017).
24. K Oh-Hashi, T Hasegawa, Y Mizutani, K Takahashi, Y Hirata, Elucidation of brefeldin a-induced ER and golgi stress responses in neuro2a cells. *Mol. cellular biochemistry* **476**, 3869–3877 (2021).
25. P Sehgal, et al., Inhibition of the sarco/endoplasmic reticulum (ER) Ca2+-ATPase by thapsigargin analogs induces cell death via ER ca2+ depletion and the unfolded protein response. *The J. biological chemistry* **292**, 19656–19673 (2017).
26. K Moncoq, CA Trieber, HS Young, The molecular basis for cyclopiazonic acid inhibition of the sarcoplasmic reticulum calcium pump. *The J. biological chemistry* **282**, 9748–9757 (2007).
27. CM Oslowski, F Urano, Measuring ER stress and the unfolded protein response using mammalian tissue culture system. *Methods enzymology* **490**, 71–92 (2011).
28. CS O'Hern, LE Silbert, AJ Liu, SR Nagel, Jamming at zero temperature and zero applied stress: The epitome of disorder. *Phys. Rev. E* **68**, 011306 (2003).
29. DS Shimamoto, M Yanagisawa, Common packing patterns for jammed particles of different power size distributions. *Phys. Rev. Res.* **5**, L012014 (2023).
30. RB Hawkes, DV Holberton, Myonemal contraction of Spirostomum I. Kinetics of contraction and relaxation. *J. Cell. Physiol.* **84**, 225–236 (1974).
